# Identification and characterization of repressive domains in *Drosophila* transcription factors

**DOI:** 10.1101/2022.08.26.505062

**Authors:** Loni Klaus, Bernardo P. de Almeida, Anna Vlasova, Filip Nemčko, Alexander Schleiffer, Katharina Bergauer, Martina Rath, Alexander Stark

## Abstract

All multicellular life relies on differential gene expression, determined by regulatory DNA elements and DNA-binding transcription factors that mediate activation and repression via cofactor recruitment. While activators have been extensively characterized, repressors are less well studied and their repressive domains (RDs) are typically unknown, as are the RDs’ properties and the co-repressors (CoRs) they recruit. Here, we develop the high-throughput next-generation-sequencing-based method Repressive-Domain (RD)-seq to systematically identify RDs in complex libraries. Screening more than 200,000 fragments covering the coding sequences of all transcription-related proteins in *Drosophila melanogaster*, we identify 195 RDs in known repressors and in proteins not previously associated with repression. Many RDs contain recurrent short peptide motifs that are required for RD function, as demonstrated by motif mutagenesis, and are conserved between fly and human. Moreover, we show that RDs which contain one of five distinct repressive motifs interact with and depend on different CoRs, including Groucho, CtBP, Sin3A or Smrter. Overall, our work constitutes an invaluable resource and advances our understanding of repressors, their sequences, and the functional impact of sequence-altering mutations.

## Introduction

Higher organisms consist of many morphologically different cell types and organs that carry out different functions in the body. Almost all cells possess the same genetic information, yet still only express certain subsets of genes. Hence, a precise regulation of gene expression must take place. The first level of regulation is transcription – the copying of DNA into an RNA transcript by RNA polymerase II. Transcription is regulated by an intricate interplay between regulatory DNA elements, transcription factor (TF) and cofactor proteins, and the RNA polymerase II machinery: TFs bind in a sequence-specific manner to regulatory DNA and recruit non-DNA-binding cofactors, i.e. co-activator or co-repressor (CoR) proteins, that mediate transcription activating or repressing cues (Reiter *et al*, 2017; Shlyueva *et al*, 2014).

TFs are modular proteins, consisting of a DNA-binding domain (DBD) and an effector domain. The effector domain can be an activating domain (AD, also called tAD) or a repressive domain (RD) and can function independently of the full-length TF (Brent & Ptashne, 1985; Lambert *et al*, 2018; Soto *et al*, 2022). Short RDs of e.g., 31 (Kruppel-RD, (Hanna-Rose *et al*, 1997)) or 55 (Engrailed-RD, (Han & Manley, 1993a)) amino acids (AA) can be sufficient to mediate repression when tethered to DNA through a heterologous DBD like the DBD of the yeast transcription factor Gal4 (Gal4-DBD; Fig. 1A). Such tethering assays have allowed the identification of RDs of various repressive TFs such as Engrailed, Snail, Cabut and others (Tolkunova *et al*, 1998; Nibu *et al*, 1998; Belacortu *et al*, 2012; Fisher *et al*, 1996; Hanna-Rose *et al*, 1997; Han & Manley, 1993b; Soto *et al*, 2022).

**Figure 1:**
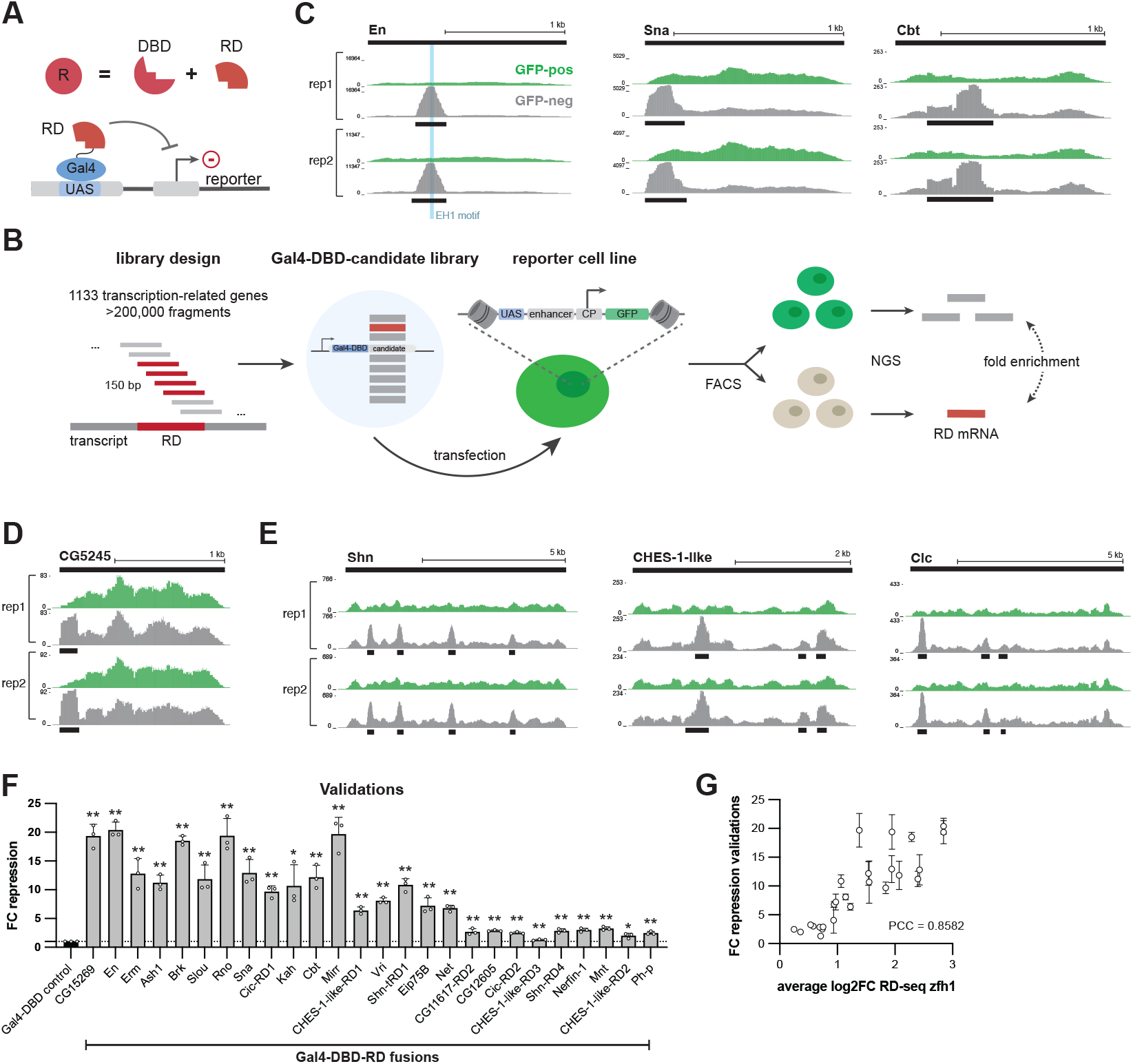
Repressive Domain-sequencing (RD-seq) identifies RDs from a comprehensive pool of candidate fragments. **A**. Repressive TFs (R) are modular and can be divided into their DNA-binding domain (DBD) and their repressive domain (RD) which is sufficient to repress a reporter when tethered to it for example via the Gal-UAS system. **B**. Schematic of the RD-seq pipeline. The candidate library consists of over 200,000 150 bp fragments tiling the coding sequences of 1133 transcription-related genes, which may contain a RD. Candidate fragments were cloned as a Gal4-DBD fusion library. Drosophila S2 cells with an integrated GFP reporter driven by a specific enhancer and core-promoter (CP) pair and with UAS sites upstream of the enhancer were transfected with the candidate library, followed by fluorescence-activated cell sorting (FACS) and next-generation sequencing (NGS). **C. – E**. UCSC genome browser tracks for two replicates of RD-seq screens with the zfh1-DSCP reporter cell line. Black bars on the top indicate the entire coding sequence of the respective factor. Shown is the normalized candidate fragment coverage from the fractions of GFP-negative and GFP-positive cells and small black bars indicating the detected RD region. **F**. Validations of RD-seq hits in comparison to the Gal4-DBD control in the zfh1-DSCP reporter cell line. Shown are the mean fold change (FC) repression values of 3 replicates and standard deviations as error bars. Significance in comparison to the Gal4-DBD control calculated with two-tailed Student T-tests is indicated above bars: * for P≤0.05, ** for P≤ 0.01. **G**. Comparison between validation FC repression values and average log2 FC in RD-seq for each RD region in the zfh1-DSCP reporter cell line. Pearson correlation coefficient (PCC) is shown.

In addition to the identification of tADs and RDs for individual TFs, pooled screening methods have been developed to systematically identify protein effector domains (Soto *et al*, 2022). Examples of such approaches include the identification of tADs within yeast, fly and human transcription factors or in random peptides (Staller *et al*, 2018; Sanborn *et al*, 2021; Staller *et al*, 2022; Erijman *et al*, 2020; Arnold *et al*, 2018; Alerasool *et al*, 2022; Ravarani *et al*, 2018) or activating and repressing domains among Pfam-annotated domains (Tycko *et al*, 2020; Alerasool *et al*, 2020). However, no systematic screen for RDs within the TF proteome of any species has been performed to date.

The sufficiency of RDs to repress transcription implies that these short domains can specifically interact with and recruit CoRs such as Groucho (Gro), CtBP and Sin3A (Jennings & Ish-Horowicz, 2008; Chinnadurai, 2002; Chaubal & Pile, 2018). Interestingly, some known RDs contain short peptide motifs which are required for RD function and are crucial for the interaction with specific CoRs. For instance, Engrailed and other repressors contain the approximately 10 AA long *engrailed-homology-1* (EH1) motif that interacts with the CoR Gro (Logan *et al*, 1992; Smith & Jaynes, 1996; Tolkunova *et al*, 1998). Similarly, the 5 AA short PxDLS motif occurs in the repressive TFs Snail and Knirps and recruits CtBP (Nibu *et al*, 1998; Quinlan *et al*, 2006). Yet, how many RDs are explained by these motifs and whether there are other peptide motifs that mediate repression and/or recruit different CoRs remains elusive.

In this study, we established *repressive-domain-sequencing* (RD-seq) to identify short 50 AA-long RDs across all annotated transcription-related proteins in *Drosophila melanogaster (Dmel)*. We recovered known and uncovered novel RDs in known repressors and in unannotated proteins. We further identified specific short peptide motifs – conserved from fly to human - and showed that RD function depends on these motifs. In addition, we used co-immunoprecipitation coupled to mass spectrometry and RNA-interference (RNAi)-mediated CoR depletion to link RD and peptide motifs to specific CoRs, revealing RD-CoR interactions and functional dependencies.

Our work provides a resource for *Drosophila* RDs as well as, the first step in building a systematic dictionary for repressors, their RDs and interacting CoRs - a valuable tool to comprehend the diverse mechanisms of transcriptional repression.

## Results

### RD-seq identifies RDs of known and novel transcriptional repressors

To systematically identify repressive protein domains (RDs), we established repressive-domain-sequencing (RD-seq), a next-generation sequencing (NGS)-based approach to identify RDs from a comprehensive pool of candidate fragments (Fig. 1B). For this purpose, we adapted the tAD-seq protocol (Arnold *et al*, 2018) and combined it with a synthetic candidate library and reporter cell lines that constitutively express GFP.

We generated a Gal4-DBD-fused candidate library consisting of over 200,000 150 bp-long DNA fragments coding for 50 AA. The candidates were designed to cover the protein-coding open-reading frames of 1,133 transcription-related *Dmel* genes in a tiled fashion with steps of 6 to 15 bp, corresponding to 2 to 5 AA (Fig. 1B, Suppl. Table 1, see library design in methods). Using CRISPR/Cas9, we created a *Dmel* S2 cell line with an integrated GFP-expressing reporter-gene cassette containing UAS sites to allow Gal4-DBD-mediated tethering of the candidates. Three days after transfection of the reporter cell line with the candidate library, we separated cells into GFP-positive and GFP-negative cells via fluorescent-activated cell sorting (FACS), followed by NGS-based quantification of the candidate mRNAs in GFP-positive and -negative cells. Since GFP-negative cells should contain candidates that repress transcription, we determined the enrichment of candidates in GFP-negative over GFP-positive cells, called RDs by their significant enrichment (p<=1×10^−5^; FC>=1.5; Fig. 1B), and for subsequent analyses only considered RDs that were detected in two of two replicates (e.g. Fig. 1C-E, see methods).

To capture different RDs, we performed RD-seq screens with two different reporter cell lines in which GFP expression was driven by distinct enhancer-promoter pairs, namely zfh1-DSCP and ent1-rps12 (Suppl. Table 2). We performed two replicates per cell line and collectively, the screens in the two cell lines resulted in a total of 195 unique RDs in 175 proteins (Suppl. Table 3). 114 of the RD-seq hits (58%) are within known or putative repressors (references in Suppl. Table 3), including the known RDs in the well-characterized repressive TFs Engrailed (En), Snail (Sna) and Cabut (Cbt) (Tolkunova *et al*, 1998; Nibu *et al*, 1998; Belacortu *et al*, 2012) (Fig. 1C). In the case of En, the peak summit of candidate enrichment coincided with the EH1 motif, known to be essential for the repressive activity (Tolkunova *et al*, 1998) (Fig. 1C blue bar in left panel). 79 RDs are in known or putative repressive TFs for which no RD had been mapped before (references in Suppl. Table 3). Moreover, we also found 81 RDs (42% of hits) in proteins that have not been implicated in repression so far, for example RDs within 18 previously uncharacterized *Dmel* proteins such as the putative Zn-finger TF CG5245 (Fig. 1D). Interestingly, some proteins have multiple RDs, for example Schnurri (Shn), for which several repressive regions have been described before (Cai & Laughon, 2009), but also CHES-1-like and Capicua (Cic) for which we identify three RDs each (Fig. 1E). Overall, RD-seq characterizes known as well as novel repressor proteins and maps RDs for both (Suppl. Table 3).

To validate RD-seq hits and assess the method’s specificity, we selected 26 of the 83 RDs that were detected in both reporter cell lines, including both strong and weak RDs from rank 1 to rank 82. We cloned a 150 bp (50 AA) fragment per RD (Suppl. Table 4), individually recruited the 26 RDs to the integrated zfh1-DSCP GFP reporter via the Gal4-DBD and assessed changes in GFP expression through flow cytometry in comparison to a control condition (Gal4-DBD alone; three independent replicates per RD and control). As a measure of the repressive strength of the RD, we calculated the fold-change (FC) repression as the median GFP signal of cells with the Gal4-DBD control versus cells with the Gal4-DBD-RD (Suppl. Fig. 1A). In the zfh1-DSCP reporter cell line, this validated all 26 hits (Student T-test P≤0.05; FC>1; Fig. 1F, Suppl. Table 4) and their repressive strengths in the validation experiments correlated well with the RD-seq enrichments (Pearson Correlation Coefficient (PCC)=0.86, Fig. 1G). Similarly, all 26 hits were validated in the ent1-rps12 reporter cell line (Suppl. Fig. 1B; Suppl. Table 4), yet the dynamic range was narrower, compressing the quantitative agreement to PCC=0.43 (Suppl. Fig. 1C). We therefore chose to use the zfh1-DSCP reporter cell line for all subsequent analyses. Overall, these results validate RD-seq as a high-throughput method to identify RDs and assess their repressive strength quantitatively.

### RDs overlap both, IDRs and DBDs, and show a preference towards N-terminal positions within a TF

Having identified and validated many RDs, we next wondered where within the TFs’ protein sequences they typically occur and analyzed the 195 RDs’ positions relative to the proteins’ N- and C-termini. As illustrated by the RDs in CG5245, Shn, CHES-1-like and Cic above (Fig. 1D, E), RDs occur at different positions within the full-length proteins. Interestingly however, they occur more frequently towards the N-termini of TFs compared to the C-termini or more intermediate positions (Fig. 2A). While the functional significance of the N-terminal positions remains unclear, the TFs’ DBDs show the opposite trend with a preference towards the C-termini of the proteins (Suppl. Fig. 2 A). These opposing trends suggest that RDs and DBDs are typically separate and non-overlapping. Indeed, only 3% of RDs overlap with DBDs (Fig. 2B), e.g. in the Hang-RD. In addition, 3% of RDs overlap with other annotated protein domains (Pfam and ProSitePatterns databases, e.g. the Parp catalytic domain in Parp-RD), 53% overlap with IDRs (MobiDB-lite database), while 41% fall into un-annotated regions (Fig. 2B and Suppl. Table 5). The large overlap of RDs and IDRs suggests an important role of these regions for repressive TFs, similar to the relevance of IDRs for activating TFs (Boija *et al*, 2018; Sabari *et al*, 2018; Chong *et al*, 2018; Brodsky *et al*, 2020; Basu *et al*, 2020). Still, many RDs don’t overlap with any known protein domains or other annotated protein features, emphasizing the need for better characterization of RDs and the protein sequence contexts in which they can function.

**Figure 2:**
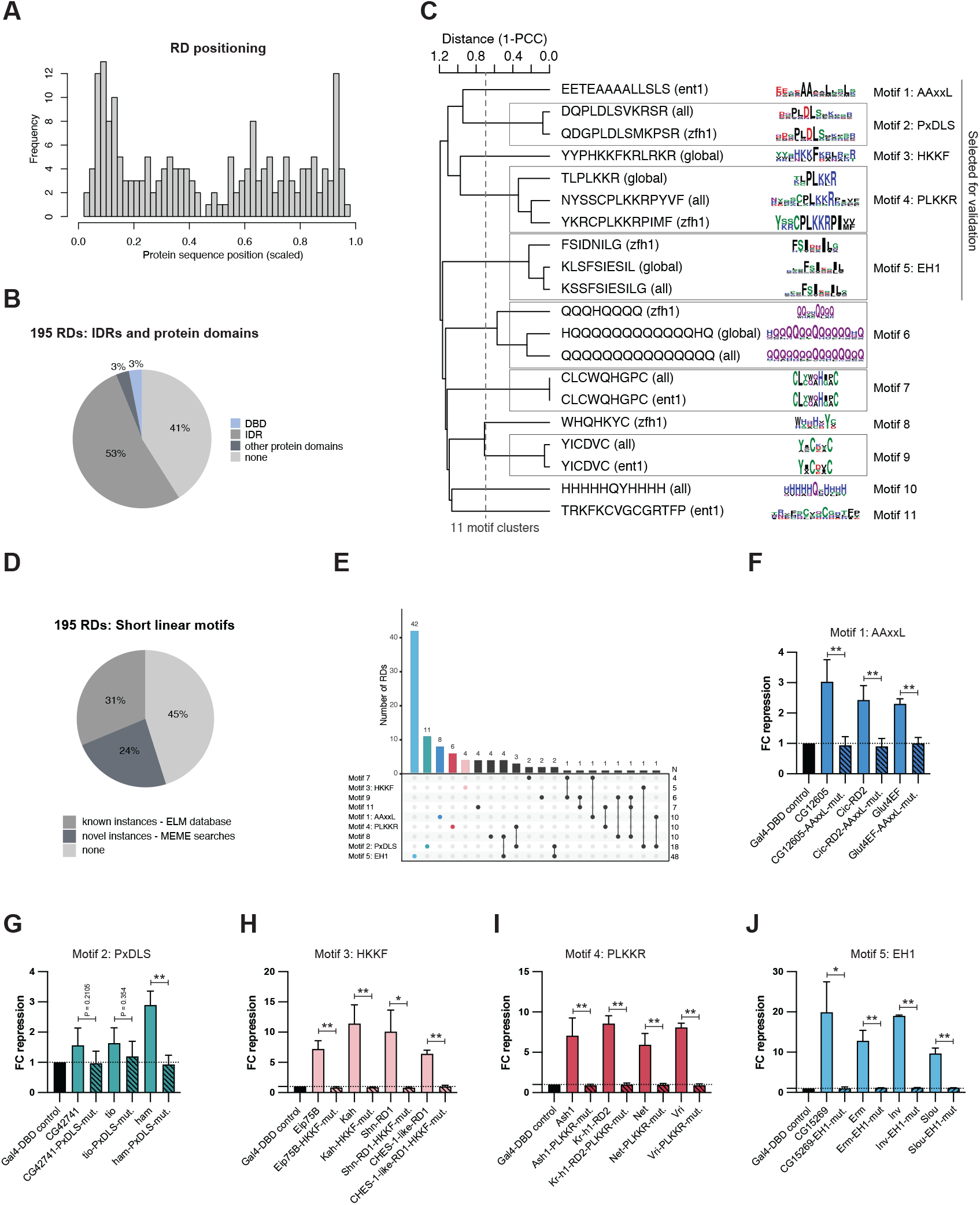
Characterization of RDs and RD dependency on short linear motifs. **A**. Frequency histogram of the position of the center of the 50 AA RD within its full-length protein for all 195 RDs. Positions are scaled over the length of the respective protein sequences. **B**. Pie chart showing the overlap between RDs and intrinsically disordered regions (IDRs) according to the MobiDB-lite database, DNA-binding domains (DBDs), and other annotated protein domains from the Pfam or ProSitePatterns databases. **C**. Hierarchical clustering of MEME *de novo* motif discovery motif hits with distinct subsets of RDs (all, global, zfh1, ent1; see methods). The tree was cut at height 0.7, resulting in 11 non-redundant distinct motifs. **D**. Pie chart showing ELM database and MEME motif instances among the 195 RDs. **E**. Number of instances of 9 MEME motifs (excluding motif 6 and 10) among the 195 RDs and co-occurrence of these motifs. N indicates the total number of RDs with a certain motif. **F. – J**. Validation results for wild type and mutated RDs in the zfh1-DSCP reporter cell line with the following conserved motifs: F. Motif 1 – AAxxL, G. Motif 2 – PxDLS, H. Motif 3 – HKKF, I. Motif 4 – PLKKR, J. Motif 5 – EH1. Shown are mean FC repression values of 3 replicates and standard deviations. Significance in comparison to the wild type RDs calculated with two-tailed Student T-tests is indicated above bars: * for P≤0.05, ** for P≤0.01, or exact P-value when not significant.

### RDs contain recurring short linear peptide motifs

As RDs can contain short peptide motifs that mediate repressor-CoR interactions (Tolkunova *et al*, 1998; Nibu *et al*, 1998), we sought to identify recurrent short peptide motifs that could explain the RDs’ repressive functions. We performed *de novo* motif discovery using MEME (Bailey *et al*, 2015) for all 195 RD-seq hits and subsets (see methods), followed by clustering of similar motifs to obtain 11 distinct short peptide motifs (Fig. 2 C, Suppl. Table 6).

Among these, we found previously annotated short-linear motifs (SLiMs) known to be important for repression and interaction with CoRs, such as AAxxL, PxDLS and EH1. The AAxxL motif resembles the Sin3A-interacting domain (SID) which recruits the CoR Sin3A (Belacortu *et al*, 2012). The PxDLS motif is known to facilitate the recruitment of CtBP (Quinlan *et al*, 2006), while the EH1 (engrailed homology 1) motif is known to mediate the interaction with the CoR Groucho (Gro) (Tolkunova *et al*, 1998; Copley, 2005). Motif 8 resembles the HCF-1 binding motif which mediates interaction with the host cell factor-1 (Hcf in *Dmel*) that has been implicated in both transcriptional activation and repression (Wysocka *et al*, 2003; Zargar & Tyagi, 2012).

In addition, motifs 7, 9, and 11 resemble zinc-finger domains from the Pfam or ProSitePatterns databases, a domain type known to mediate DNA-binding, protein-protein interactions (reviewed in Brayer & Segal, 2008), but also transcriptional repression (Tapia-Ramírez *et al*, 1997; Lee *et al*, 2005). Two additional motifs were of low sequence complexity with multiple glutamate (motif 6) or histidine residues (motif 10), which have been observed in activating and repressing TFs (Ramazzotti *et al*, 2012; Atanesyan *et al*, 2012; Salichs *et al*, 2009). Hence, to avoid studying compositional biases of transcriptional regulators in general, we excluded the Q and H repeat motifs from further analysis, and instead focused on the other 9 MEME motifs.

We also found two novel, previously unannotated motifs, motifs 3 and 4, that we termed PLKKR and HKKF, respectively. The two motifs are potentially novel SLiMs and both are positively charged, consistent with the positive charges in recently identified repressive domains (Tycko *et al*, 2020).

We next mapped the positions of all instances of the 9 main motifs within the 195 RDs (Suppl. Table 6), as well as motif instances from the ELM database (Suppl. Table 7). Of all 195 RDs, 55% contain at least one instance of these motif types, of which 24% could only be identified with the *de novo* defined motifs (Fig. 2D). For the AAxxL motif, we find multiple novel instances, e.g. in the RDs of Glut4EF, CG12605 and Cic (Suppl. Table 6). Interestingly, EH1 was the most abundant motif, present in 48 different RDs (Fig. 2E). This large group of EH1 motif-containing RDs is the main driver of the positional bias of RDs towards the N-termini of the full-length TFs (Fig. 2A and Suppl. Fig. 2B). Moreover, some RDs contain combinations of peptide motifs, such as the RDs of Sna, Esg and Wor that all contain both, the PxDLS and the PLKKR motif (Fig. 2E).

### Short peptide motifs are required for RD function

Next, we assessed the necessity of five of the known and novel peptide motifs for the repressive activity of RDs by mutating the motifs to Alanine residues. We selected motif types 1 through 5 (i.e. AAxxL, PxDLS, HKKF, PLKKR and EH1) and mutated between three and four different RDs per motif type (Suppl. Table 4 with all AA sequences). We first confirmed that the mutated RDs were still expressed to equal or higher levels compared to the wild type RDs (Suppl. Fig. 2C), such that changes in the repressive activity are not caused by impaired protein stability. For all instances of all motif types, we observed a loss of repressive activity upon motif mutation, rendering the mutated RD variants as ineffective as a Gal4-DBD control (Fig. 2F-J, Suppl. Table 4). For two weaker PxDLS motif containing RDs the loss of repression was not significant but still noticeable (Fig. 2G). The results so far reveal known as well as novel short peptide motifs and show that RDs rely on such motifs to repress transcription.

### RDs with different peptide motifs bind distinct co-repressors

Some of the recurrent peptide motifs that are essential for RD function have been described previously to facilitate the interaction between repressors and CoRs (Tolkunova *et al*, 1998; Nibu *et al*, 1998; Belacortu *et al*, 2012). After validating the motifs’ necessity for RD function, we wanted to explore their mechanism of action, specifically the CoR proteins they might recruit. To determine the interactors of RDs with different motifs, we performed immunoprecipitations of RDs followed by quantitative mass spectrometry (IP-MS). IPs were performed using an anti-FLAG antibody and nuclear lysate of *Drosophila* S2 cells overexpressing 3xFLAG-Gal4-DBD-tagged RDs with a specific peptide motif or 3xFLAG-Gal4-DBD as negative control. To ensure that each motif is in the sequence context of a functional RD, while also ensuring that binding partners of the motif rather than any individual RD are characterized, we performed immunoprecipitation experiments with pools of several RDs that share the motif of interest. We excluded EH1 motif containing RDs, since this motif and its interaction with Gro has already been studied extensively (Tolkunova *et al*, 1998; Jennings *et al*, 2006; Copley, 2005).

As expected, RDs containing the PxDLS motif enriched for the CoR CtBP (Nibu *et al*, 1998) (Fig. 3A) and AAxxL motif-containing RDs enriched for the Sin3A CoR complex members Sin3A, HDAC1 and CG14220 (Belacortu *et al*, 2012) (Fig. 3B). RDs with the PLKKR motif interacted with four subunits of the Smrter CoR complex (orthologous to human NCoR/SMRT), namely Smr, CG17002, Ebi and HDAC3 (Fig. 3C). While the PLKKR motif has not been described as a CoR-interacting SLiM, our IP-MS results are consistent with two studies describing the interaction of *Dmel* Snail with the Smrter subunit Ebi through a YxxCPLKKRP sequence (Qi *et al*, 2008) and the human MeCP2 protein with the Ebi ortholog TBLR1 through an extended domain that contains a PIKKR sequence (Lyst *et al*, 2013; Kruusvee *et al*, 2017). Our data suggests that the PLKKR motif is a recurrent SLiM utilized by various repressive TFs which likely mediate transcriptional repression through the Smrter CoR complex.

**Figure 3:**
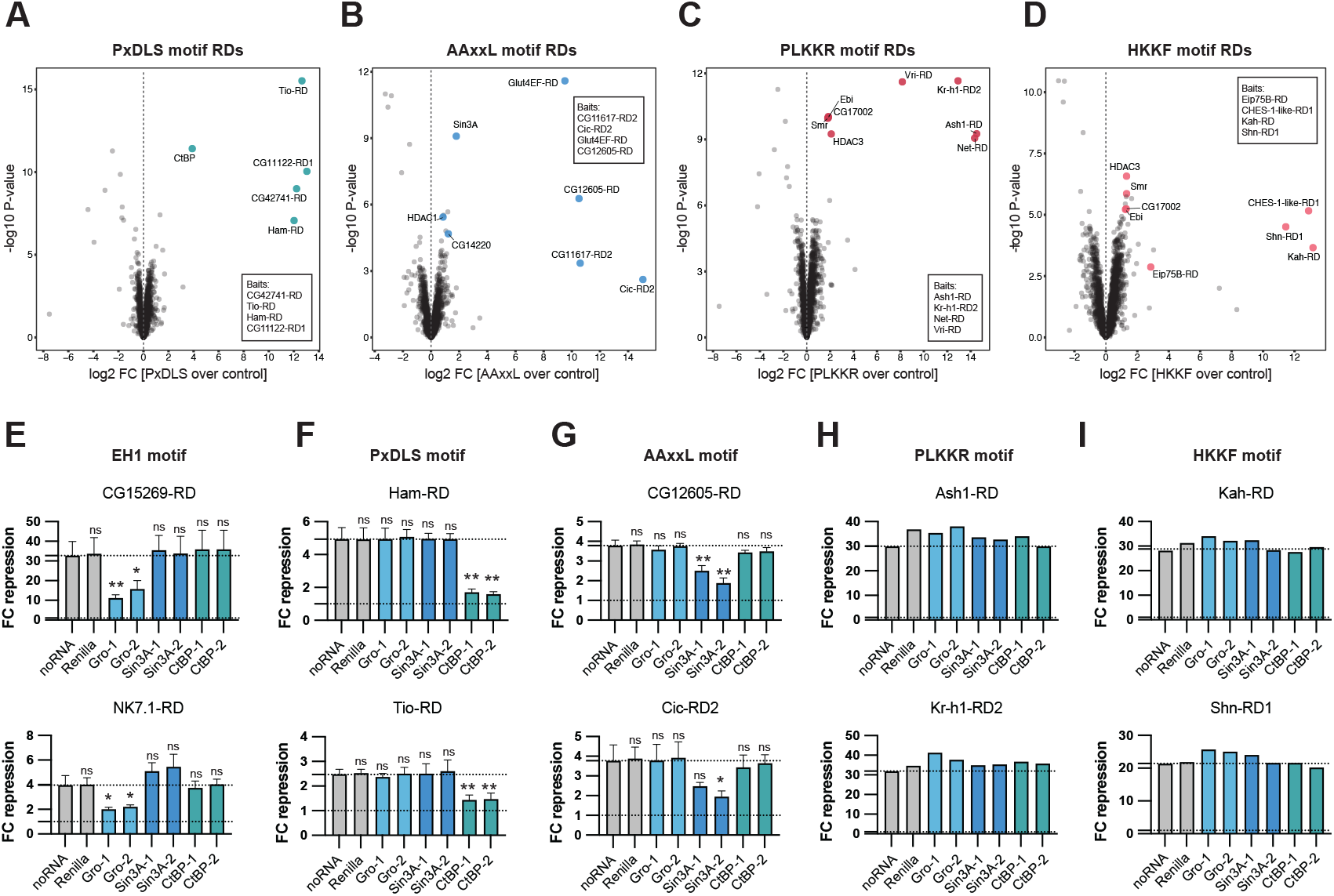
RD – CoR interactions and dependencies. **A. – D**. Results of immunoprecipitations followed by mass spectrometry (IP-MS) for pools of RDs with specific repressive motifs: A. PxDLS, B. AAxxL, C. PLKKR, D. HKKF. Shown are volcano plots with the log2FC over control on the x-axis and the -log10 P-value on the y-axis. The FLAG-Gal4-DBD-tagged RDs used as bait for the IPs are indicated in boxes. **E. – I**. Validations of RDs upon RNAi-mediated depletion of CoRs in the zfh1-DSCP reporter cell line. Each CoR was targeted for depletion with 2 different dsRNA constructs. A dsRNA targeting Renilla and a condition without any dsRNA (noRNA) were used as controls. Shown are means of FC repression values of 3 replicates with standard deviations (E.-G.) or the FC repression value of 1 replicate (H., I.). The repressive motif contained in the tested RD is indicated above the panels. For E.-G. significance in comparison to the noRNA control calculated with two-tailed Student T-tests is indicated above bars: * for P≤0.05, ** for P≤0.01, or not significant (ns) for P>0.05.

Interestingly, RDs with the HKKF motif also enriched for the Smrter complex (Fig. 3D), consistent with a report that the repressive TF Shn interacted with Smrter via a NISRYLHKKFKRLASTTEVDS sequence (Cai & Laughon, 2009). This sequence not only contains a HKKF motif but also coincides with the first of four RDs we find within Shn (Fig. 1E and Fig. 2H). Hence, similar to PLKKR, the HKKF motif is a SLiM likely utilized by various repressors to interact with the Smrter complex.

### RDs with distinct peptide motifs depend on different co-repressors

We set out to corroborate the results of the IP-MS experiments by assessing CoR requirements for RD function. We designed dsRNAs for the RNAi-mediated depletion of four different CoRs by dsRNA transfection in *Drosophila* S2 cells. RT-qPCRs showed the successful depletion of Gro, CtBP and Sin3A mRNAs through treatment with two distinct dsRNAs each (Suppl. Fig. 3A). However, we could not sufficiently strongly deplete the transcripts of Smr or Ebi despite the use of two different dsRNA constructs each and therefore could not follow up on the dependency of RDs on the Smrter complex.

RNAi-mediated CoR depletion revealed that EH1 motif-containing RDs specifically depended on Gro but not on CtBP or Sin3A (Fig. 3E). In contrast, PxDLS motif-containing RDs depended on CtBP but not Gro or Sin3A (Fig. 3F) and AAxxL motif-containing RDs required Sin3A but not the other two CoRs (Fig. 3G). Each of these dependencies was consistent with literature reports (Tolkunova *et al*, 1998; Jennings *et al*, 2006; Copley, 2005) or the IP-MS results for the different motifs (Fig. 3A, B). Interestingly, RDs with PLKKR or HKKF motifs maintained their repressive function in the absence of each of these 3 CoRs (Fig. 3H, I), which indicates that these motifs are independent of Gro, CtBP and Sin3A, in line with their likely dependence on the Smrter CoR complex.

Overall, our experiments suggest that repressors mediate repression through short, conserved peptide motifs which are required for the interaction with certain CoRs. Interestingly, some repressors contain multiple RDs that recruit different types of CoRs, for example Schnurri with RDs for Gro, Sin3A and Smrter, and CHES-1-like with RDs for Smrter and Sin3A (Fig. 1E, Suppl. Fig. 3B). There are also cases in which RDs contain two distinct peptide motifs, such as PLKKR and PxDLS within the RD of Snail (Fig. 2E, Suppl. Fig.3B). Investigating different Snail-RD mutants showed that both motifs contribute to the RD’s repressive activity (Suppl. Fig. 3C): mutating the PxDLS motif alone does not impair RD function and while mutating the PLKKR motif deceases RD function, only the simultaneous mutation of both motifs abolishes it. Consistently, the RD of Sna remains functional when CtBP is depleted by RNAi (Suppl. Fig. 3D), presumably because it is still able to recruit the Smrter complex via its PLKKR motif. These observations of proteins with multiple RDs and likely different interacting CoRs have interesting implications for how even single transcriptional repressors could act in different ways to achieve gene silencing.

### Fly RD motifs are conserved across species and predict human repressors

Some of the RD motifs and their interactions with CoRs are known to be conserved across species as distant as flies and mammals. This includes the EH1, PxDLS and AAxxL motifs and their interaction with the human orthologs of Gro, CtBP and Sin3A, which have been described for individual human proteins (Logan *et al*, 1992; Quinlan *et al*, 2006; Belacortu *et al*, 2012).

To illustrate the deep conservation of individual instances of these motifs, we created sequence alignments for repressive TFs from *Dmel* containing repressive peptide motifs and the TFs’ orthologs in different species over a wide phylogenetic range (Fig. 4A, B; Suppl. Fig. 4A, B). The alignment of Eip93F and its orthologs illustrates the conservation of the PxDLS motif from insects to mammals (Fig. 4A), as does the alignment of Mid and its orthologs for the EH1 motif (Suppl. Fig. 4A). The alignment of Glut4EF containing the AAxxL motif shows the strong conservation not only of the core AAxxL motif but also of the flanking sequences, suggesting that this motif might in fact be longer (Suppl. Fig. 4B). Lastly, also the PLKKR motif within Vri is strongly conserved in Vri’s orthologs (Fig. 4B).

**Figure 4:**
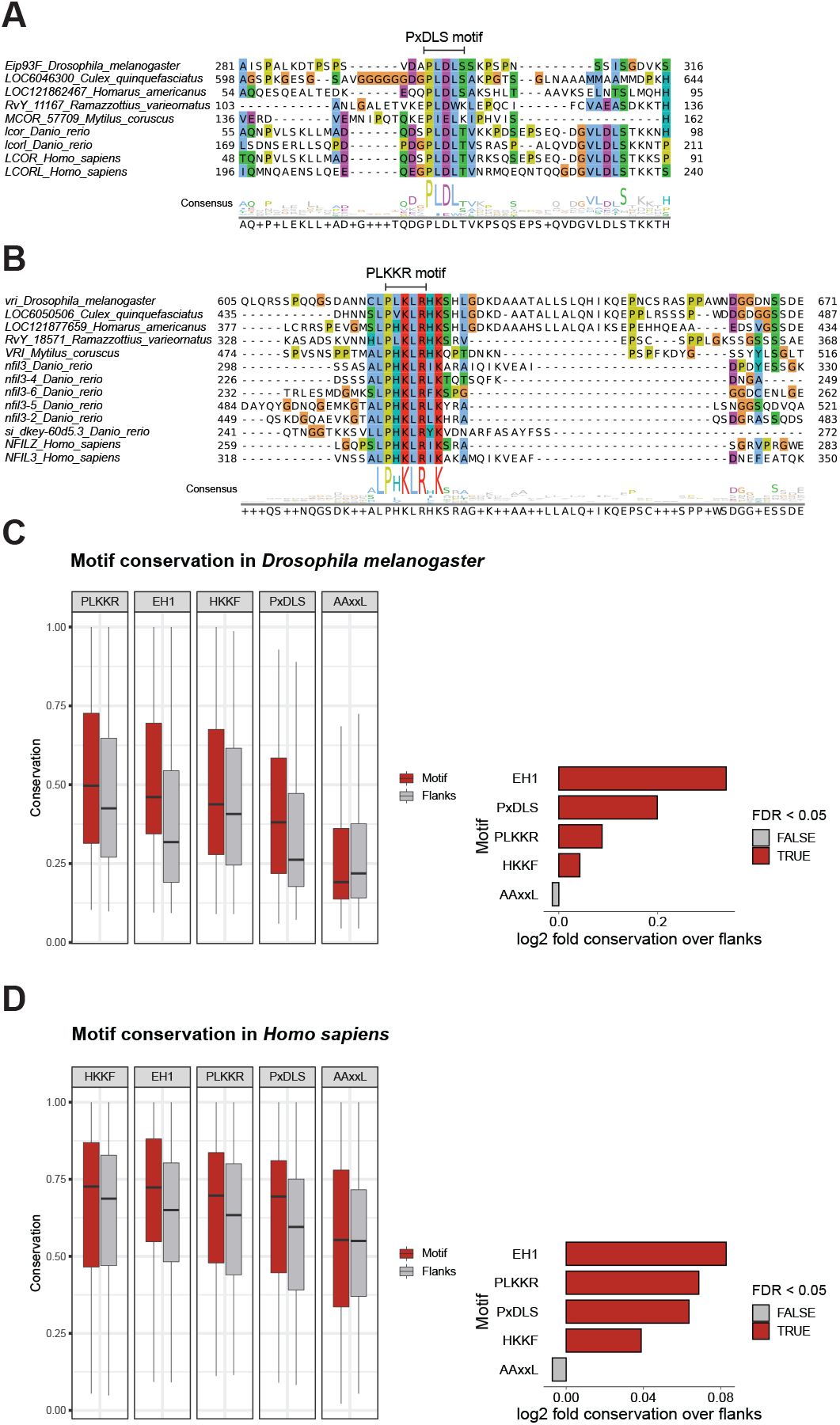
RDs and repressive peptide motifs are conserved across species. **A. B**. Sequence alignments for a region of *Dmel* TFs (A) Eip93F containing the PxDLS motif or (B) Vri containing the PLKKR motif and the respective orthologous sequences from different species. Numbers on the left and right indicate the range of amino acids shown referring to the full-length proteins. Consensus sequences are indicated on the bottom. **C. D**. Conservation of repressive motifs over their flanking regions for motif instances from all (C) *Dmel* and (D) human transcription-related genes. Left: Box plots with average conservation scores of motif instances (red) and respective flanking regions (grey) for each motif type. Right: Bar plot with log2 fold conservation between motif instances and their flanking regions per motif type. Motifs are colored by significance: two-sided Wilcoxon rank-sum test FDR-corrected p-value <0.05.

If instances of the repressive motifs EH1, AAxxL, PxDLS, PLKKR and HKKF were indeed functional in fly and human TFs, they would on average be more highly conserved than expected in closely related insect or vertebrate species, respectively. This reasoning has previously been applied to short microRNA-binding-site sequences in flies and mammals (Brennecke *et al*, 2005; Lewis *et al*, 2005) or TF binding sites (e.g. Stark *et al*, 2007) and benefits from the better alignability of sequences between closely related species rather than distal ones. Following this reasoning, we created multiple protein-sequence alignments of *Dmel* and human transcription-related proteins within insect and vertebrate orthogroups, respectively. We next calculated conservation scores for each AA position of these transcription-related proteins from *Dmel* (same as the proteins in RD-seq library, Suppl. Table 1) and human (based on Lambert *et al*, 2018; Vaquerizas *et al*, 2009) (Suppl. Table 8) and assessed the conservation of the 5 different peptide motifs (Fig. 4C, D) compared to immediately flanking sequences. In both insects and vertebrates, we observed significantly higher conservation of the EH1, PLKKR, PxDLS and HKKF motifs in comparison to their flanking regions. The AAxxL motif, however, was not significantly more highly conserved than its flanks for both, insects, and vertebrates. Therefore, even though the AAxxL motif validated experimentally and AAxxL-motif-containing RDs interacted with Sin3A and depended on Sin3A (Fig. 2F, 3B, 3G), its function is not reflected by increased conservation compared to its flanks. This might be due to the motif being longer and extending into the flanks, as exemplified by the alignment of Glut4EF (Suppl. Fig. 4B). The increased conservation of the RD motifs compared to the flanking sequences suggests that at least EH1, PxDLS, PLKKR and HKKF are under purifying selection in both insects and vertebrates and thus likely functionally relevant (e.g. Lewis *et al*, 2005; Brennecke *et al*, 2005).

Among the 2754 human transcription-related proteins (based on Lambert *et al*, 2018; Vaquerizas *et al*, 2009) that contained repressor motifs (Suppl. Tables 8 & 9) were indeed many known repressors: for example among the 30 highest scoring PxDLS motif matches we found 19 proteins known to repress transcription through CtBP, for example MECOM (also EVI1), ZFPM1 and PRDM16 (Izutsu *et al*, 2001; Katz *et al*, 2002; Kajimura *et al*, 2008; additional references in Suppl. Table 9). The highest scoring PLKKR matches include MeCP2, known to contain a PIKKR sequence and to interact with NCoR/SMRT (Kruusvee *et al*, 2017), and other proteins that have been implicated in repression but not been associated with any CoR, such as NSD2 and ASH1L (Nimura *et al*, 2009; Tanaka *et al*, 2011) (Suppl. Table 9). Similar to the situation in *Dmel*, (see Suppl Fig. 3B) some human proteins like BCL3 contain both the PxDLS and PLKKR motifs, suggesting that they recruit both, the CtBP and the NCoR/SMRT CoR complexes.

These analyses not only highlight the deep evolutionary conservation of repressive peptide motifs but also provide both, an annotation of human repressive TFs that contain such motifs and a resource to study human TF sequences and assess the potential functional impact of mutations in these proteins.

## Discussion

Transcriptional activation and repression are both crucial for gene regulatory programs in different cell types and under changing environmental conditions. Yet, while transcriptional activators and trans-activating domains (tADs) have been studied extensively (Ravarani *et al*, 2018; Staller *et al*, 2018; Arnold *et al*, 2018; Erijman *et al*, 2020; Sanborn *et al*, 2021; Alerasool *et al*, 2022; Staller *et al*, 2022), our knowledge on transcriptional repressors, their RDs and interacting CoRs remained limited. Here, we developed the high-throughput assay RD-seq, to systematically map RDs throughout the sequences of all transcription-related proteins in *Dmel* (Fig. 1). This identified 195 unique RDs in known repressors and proteins that have not been implicated in repression, providing the first comprehensive screen for RDs and a resource for RD – CoR associations.

We find that RDs contain short recurring peptide motifs required for the RDs’ repressive functions (Fig. 2), and these motifs recruit specific CoRs as demonstrated by IP-MS and functional RD-CoR dependencies (Fig. 3). These include known examples (Tolkunova *et al*, 1998; Nibu *et al*, 1998; Belacortu *et al*, 2012) such as the well-established EH1-Gro and PxDLS-CtBP interactions (Tolkunova *et al*, 1998; Jennings *et al*, 2006; Copley, 2005; Nibu *et al*, 1998; Ryu & Arnosti, 2003) and the less well-studied interaction of AAxxL and Sin3A (Zhang *et al*, 2001; Belacortu *et al*, 2012). Furthermore, our study reveals two new recurrent SLiMs, PLKKR and HKKF, found in RDs that bind the Smrter CoR complex (Fig. 2, Fig. 3). This finding is consistent with two studies reporting the interaction between extended fly or human protein domains with the Smrter or NCoR/SMRT complex, respectively (Qi *et al*, 2008; Kruusvee *et al*, 2017; Cai & Laughon, 2009). Our results refine these studies to pinpoint PLKKR- and HKKF-like motifs in these domains. Indeed, point mutations within MeCP2 that lead to the Rett syndrome (Lyst *et al*, 2013; Kruusvee *et al*, 2017) map to the PIKKR motif, highlighting the importance and potential disease-association of RDs.

The lack of systematic annotations of RDs in fly TFs makes it difficult to evaluate the specificity and sensitivity of RD-seq against an independent benchmark dataset. However, the candidate library contained fragments covering 438 TFs whose regulatory activity was assessed in a previous study (Stampfel *et al*, 2015). We found RDs in 79 of these TFs, of which 50 (63%) are repressors, which increases to 61 (77%) for TFs that are at least weakly repressive and 73 (92%) for TFs that are not activators (see methods). In addition, we recover a variety of RDs that have been mapped in studies on individual repressive TFs (e.g. Tolkunova *et al*, 1998; Hemavathy *et al*, 2004; Cai & Laughon, 2009) (more references in Suppl. Table 3). These results suggest that RD-seq is highly specific, consistent with the validation rate of 26 out of 26 RDs (Fig. 1F). Of the 156 repressive TFs derived from Stampfel *et al*. (2015), we found RDs for 50 (32%), and for the 43 strongly repressive TFs, we found RDs in 22 (51%). The recovery of RDs in these sets of TFs increased to 66 (42%) and 27 (63%), respectively, when calling RDs with a more lenient threshold in RD-seq (see methods). The remaining repressors might require specific cellular or regulatory contexts to function or contain RDs that are too weak to be detected by RD-seq, are bipartite, and/or are longer than the 50 AA fragments we screened.

A majority of the identified RDs (55%) contain recurrent motifs that might explain their CoR interactions and functions. The remaining 45% of RDs did not contain any of these SLiMs, suggesting that they function via rare motifs shared between only very few RDs (precluding the motifs’ discovery by statistical over-representation) or by entirely different means. Some RDs may use different motifs to recruit the same CoR, as has been described for the EH1 and the WRPW motifs that both recruit Gro (Tolkunova *et al*, 1998; Fisher *et al*, 1996). Other RDs may utilize entirely different sets of CoRs than the ones found and studied here. Which kind of repressive mechanisms different repressor-CoR pairs utilize remains an open question for future research. Interestingly, we found several examples of repressors with multiple RDs harboring distinct repressive motifs and likely recruiting different CoRs (Fig. 1E, Suppl. Fig. 3B). Such motif-based modularity could allow for additive functions of transcriptional repressors.

Strikingly, the properties of RDs differ remarkably from those of tADs (Brent & Ptashne, 1985; Arnold *et al*, 2018). While many RDs contain conserved repressive motifs (Fig. 4) that bind specific CoRs (Fig. 3), tADs don’t share recurrent motifs, are typically poorly conserved and difficult to predict (Erijman *et al*, 2020; Sanborn *et al*, 2021; Soto *et al*, 2022; Erkina & Erkine, 2016). Moreover, tADs have been described to show rather fuzzy and weak binding of their cofactors (Erijman *et al*, 2020; Sanborn *et al*, 2021) with variable binding interfaces (Sanborn *et al*, 2021). Yet, the presence of recurrent conserved repressive motifs in RDs suggests well defined RD-CoR interaction interfaces, which for some examples have indeed been described by structural studies (Jennings *et al*, 2006; Nardini, 2003; He *et al*, 2021). These differences in RD and tAD characteristics are interesting because they indicate that transcriptional activation and repression utilize different biochemical mechanisms and principles to cause opposite effects on gene expression.

Notably, the RD properties uncovered in *Dmel* are shared with human repressors: Repressive motifs found in *Dmel* are deeply conserved throughout evolution (Fig. 4), and the annotation of RDs through such motifs poses a valuable resource for studying RDs and the impact of RD mutations, for example in disease contexts. Understanding RDs and their interacting CoRs is particularly important at a time when interests are increasingly shifting from studying transcriptional activation towards the actors and mechanisms of transcriptional repression.

## Acknowledgements

We thank Lorena Hofbauer, Jelle Jacobs and other members of the Stark group for discussions, Lisa-Marie Pleyer (IMP) for help with GEO submission, and Filip Nemčko and Lorena Hofbauer (IMP) for comments on the manuscript. We thank Michaela Pagani (IMP) for support with experiments. Next generation sequencing was done at the Vienna Biocenter Core Facilities GmbH (VBCF) Next-Generation Sequencing Unit (http://vbcf.ac.at); FACS and flow cytometry at the BioOptics facility of the IMP/IMBA/GMI. We thank the mass spectrometry unit at IMP/IMBA/GMI for their help with IP-MS experiments. We thank the Brennecke Lab (IMBA) for sharing materials and for discussions. Loni Klaus is a recipient of a DOC Fellowship of the Austrian Academy of Sciences at the Research Institute of Molecular Pathology. Filip Nemčko is supported by a Boehringer Ingelheim Fonds PhD fellowship. Research in the Stark group is supported by the Austrian Science Fund (FWF, P29613-B28). Basic research at the IMP is supported by Boehringer Ingelheim GmbH and the Austrian Research Promotion Agency (FFG). For the purpose of Open Access, the author has applied a CC-BY-NC-ND 4.0 International license to this pre-print.

## Author Contributions

LK and ASt conceptualized the study. LK performed most of the experimental work and data analyses and prepared the figures. BPdA performed bioinformatic analysis on RD characteristics and conservations of repressive motifs over their flanking sequences. AV performed NGS data analysis. ASc prepared evolutionary sequence alignments and calculated conservation scores for fly and human proteins. FN, KB, and MR helped with experiments. LK and ASt wrote the manuscript with help from BPdA. ASt supervised the study.

## supplementary Figures

**Suppl. Figure 1:**
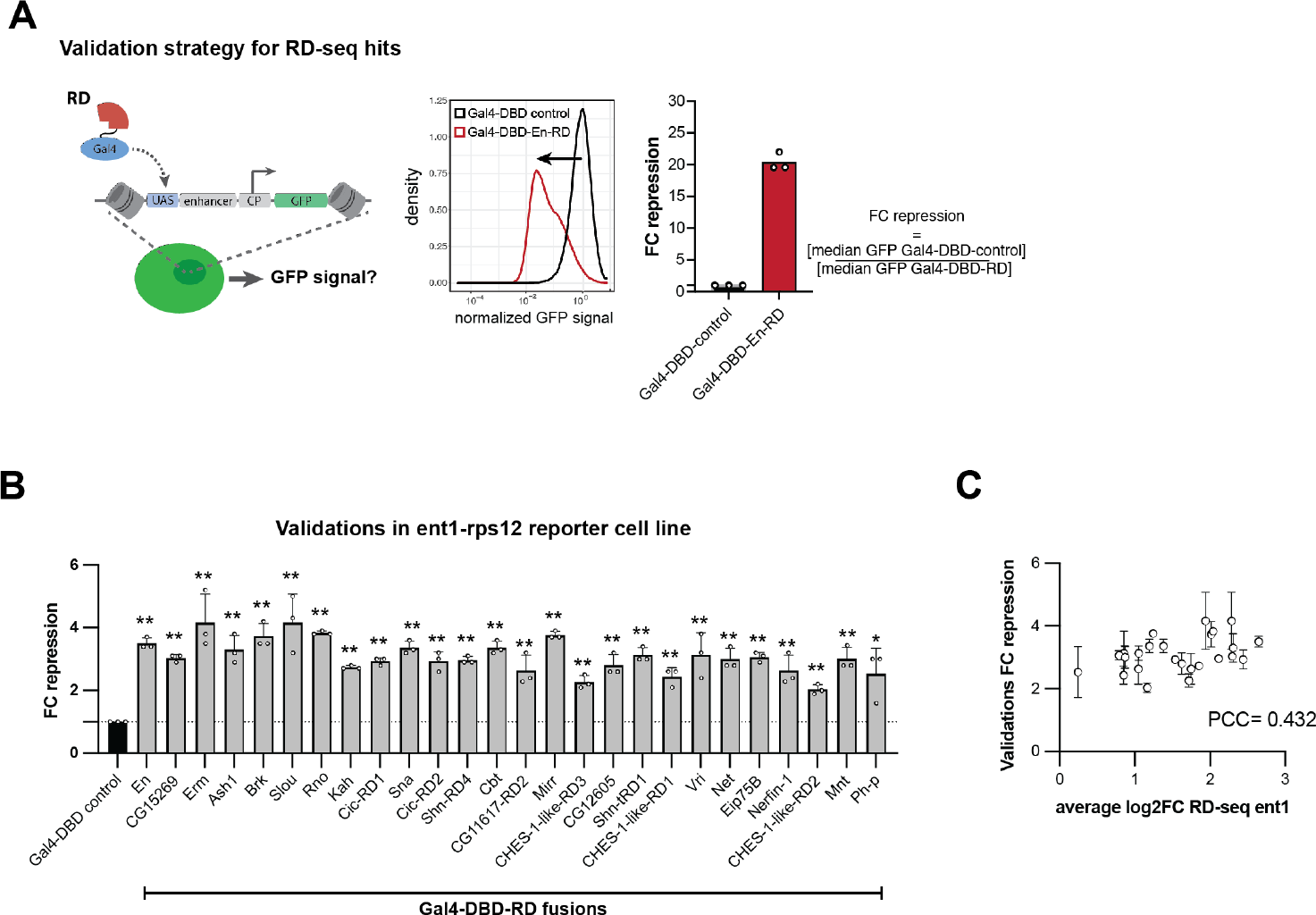
Validations of RD-seq hits. **A**. Validation strategy for RD-seq hits. The reporter cell line is transfected with either the Gal4-DBD-fused RD or a Gal4-DBD construct as a control, followed by assessment of the GFP-signal by flow cytometry (left). The density distribution shows the normalized GFP signal of cells expressing either the Gal4-DBD control or the Gal4-DBD-En-RD construct (middle). The fold change (FC) repression, i.e. the RD strength, is calculated as the ratio of the median GFP intensity of the Gal4-DBD control and the Gal4-DBD-RD condition. Shown is the mean of 3 replicates and individual values for the Gal4-DBD control and the Gal4-DBD-En-RD condition (right). **B**. Validations of RD-seq hits in comparison to the Gal4-DBD control in the ent1-rps12 reporter cell line. Shown are the mean Fold change (FC) repression values from 3 replicates and standard deviations as error bars. Significance in comparison to the Gal4-DBD control calculated with two-tailed Student T-tests is indicated above bars: * for P≤0.05, ** for P≤0.01. **C**. Comparison between validation FC repression values and average log2 FC in RD-seq for each RD region in the ent1-rps12 reporter cell line. Pearson correlation coefficient (PCC) is shown.

**Suppl. Figure 2:**
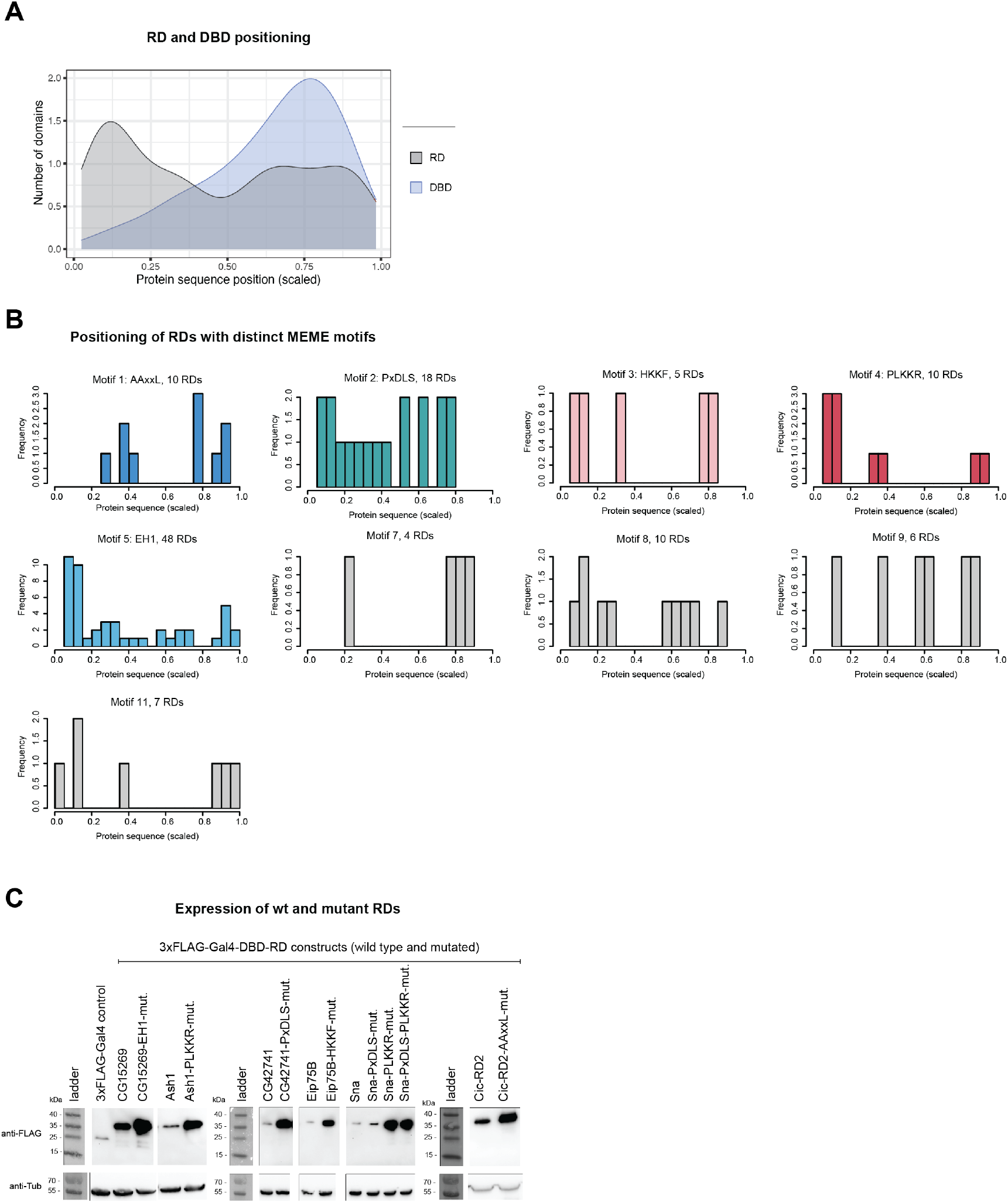
RD and DBD positioning and expression of mutated RDs. **A**. Positioning of RDs and DBDs. Density distribution of the position of the center of the 50 AA RD or the DBD regions within their full-length protein. Positions are scaled over the length of the respective protein sequences. **B**. Positioning of RDs with distinct motifs from MEME *de novo* motif searches. Shown are frequency histograms of the position of the center of the 50 AA RD within its full-length protein for all RDs containing each motif type. Positions are scaled over the length of the respective protein sequences. **C**. Western blots for FLAG-Gal4-DBD-tagged wild type and motif mutant RDs expressed in the zfh1-DSCP reporter cell line. Blots were probed with an anti-Tubulin antibody as loading control.

**Suppl. Figure 3:**
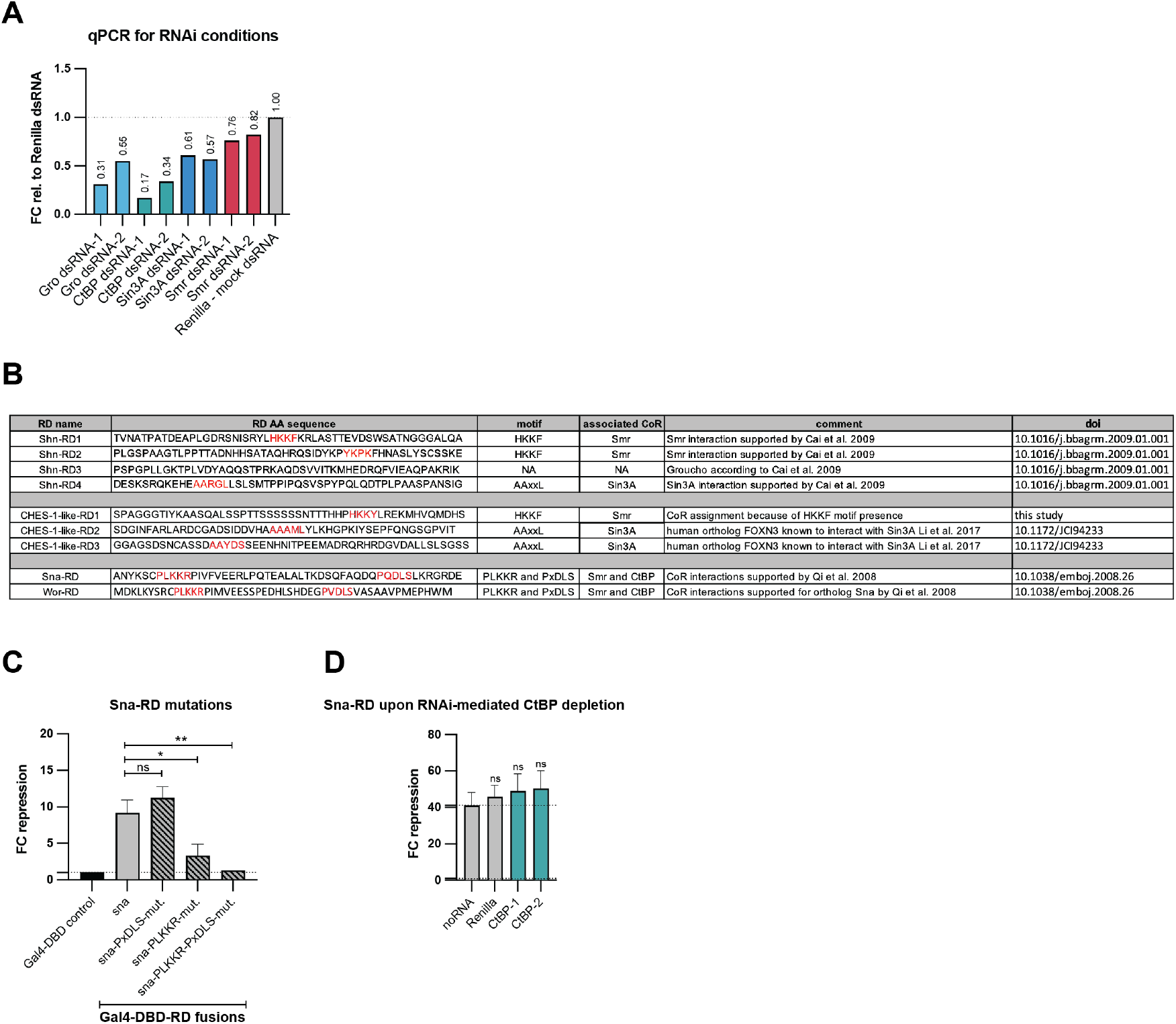
RNAi-mediated co-repressor depletion and repressors with multiple RDs and repressive motifs. **A**. Assessment of depletion of CoR mRNA with RNAi through reverse transcription quantitative PCR (RT-qPCR). Each CoR was targeted with 2 different dsRNA constructs. A dsRNA targeting Renilla was used as a negative control. Shown is the fold change (FC) relative to the control condition for one replicate each calculated with the Delta-Delta Ct Method. **B**. Repressors with multiple RDs and RDs with multiple repressive motifs. Shown are RD sequences, presence of repressive motifs, their associated interacting CoRs and literature references. **C**. Validation of wild type and mutant Sna-RD in the zfh1-DSCP reporter cell line. Shown are mean FC repression values of 3 replicates and standard deviations. Significance in comparison to the wild type RD calculated with two-tailed Student T-tests is indicated above bars: * for P≤0.05, ** for P≤0.01, not significant (ns) for P>0.05. **D**. Validation of Sna-RD upon RNAi-mediated depletion of CtBP in the zfh1-DSCP reporter cell line. CtBP was targeted for depletion with 2 different dsRNA constructs. A dsRNA targeting Renilla and a condition without any dsRNA added (noRNA) were used as controls. Shown are means of FC repression values of 3 replicates with standard deviations. Significance in comparison to the noRNA control calculated with two-tailed Student T-tests is indicated above bars: * for P≤0.05, ** for P≤0.01, or not significant (ns) for P>0.05.

**Suppl. Figure 4:**
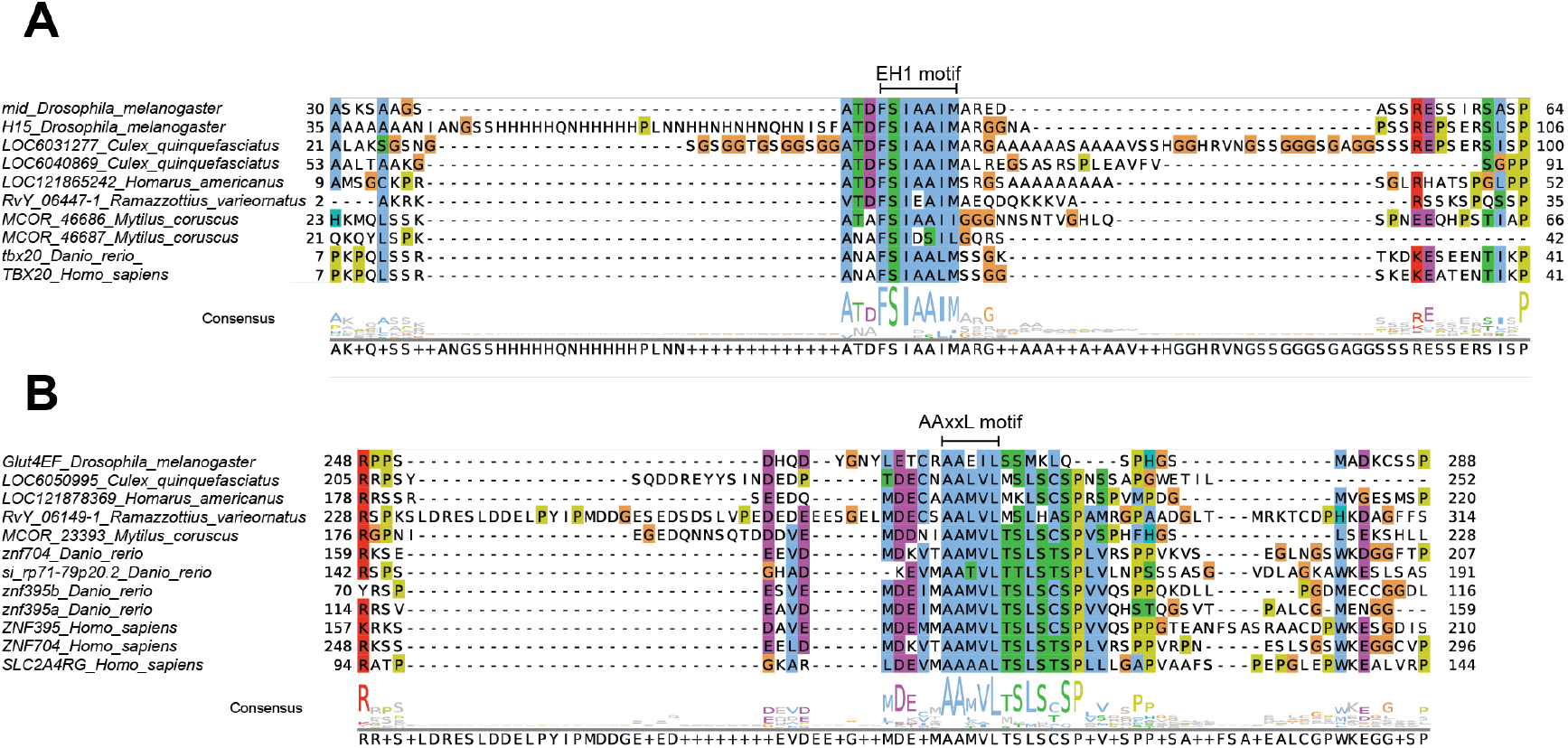
Sequence alignments of RDs. **A. B**. Sequence alignment for a region of *Dmel* TFs (A) mid and H15 containing the EH1 motif or (B) Glut4EF containing the AAxxL motif and the respective orthologous sequences from different species. Numbers on the left and right indicate the range of amino acids shown referring to the full-length proteins. Consensus sequences are indicated at the bottom.

## supplementary Tables

supplementary Tables are available upon request.

## Methods

### RD candidate expression plasmids

#### RD-seq plasmid backbone

The plasmid backbone for the RD-seq candidate library was derived from ptAD-seq-ubi63E-Gal4-DBD (Arnold *et al*, 2018) by replacing the ubi63E enhancer with the zfh1 enhancer (from pGL3_zfh1_CP-candidate_luc+; Addgene 86391) in between the KpnI (Thermo) and BglII (Thermo) restriction sites (Suppl. Table 2, RD-seq backbone: zfh1-DSCP-Gal4-DBD, primers in Suppl. Table 10). The plasmid contains the Gal4-DBD followed by a poly-glycine linker upstream of the candidate library insertion site, which consists of the *ccdB* suicide gene flanked by homology arms, which is followed by three stop codons. For details on how candidate fragments were integrated into the RD-seq backbone see *Candidate tiling library design and cloning*.

#### Validation plasmid backbone

For validation experiments we introduced the fluorescent protein EBFP2 (source: Addgene 54665) driven by the dpse enhancer and the CG13116 promoter in the RD-seq plasmid backbone to be able to gate for transfected cells in flow cytometry (Suppl. Table 2, validation backbone: zfh1-DSCP-Gal4-dpse-EBFP2). An oligonucleotide with the EBFP2 gene, a stop codon and the SV40 poly-A site synthesized by IDT (Suppl. Table 10: EBFP2-stop-polyA) was amplified with primers including overhangs for Gibson cloning (Suppl. Table 10: EGFP2_fw and _rv). The dpse enhancer and the CG13116 promoter were amplified from pAGW-dpse-GAL4-DBD (Addgene 125153) with primers including overhangs for Gibson cloning (Suppl. Table 10: dpse-CG13116-promoter_fw and _rv). Using Gibson assembly (NEB), both fragments were integrated into the LguI-linearized (Thermo) RD-seq plasmid.

#### FLAG-tag plasmid backbone

For testing the expression of mutated RDs in western blots and for IP-MS experiments the validation construct was further modified by introducing a sequence containing the 3xFLAG-tag and a short Gly-Ser linker upstream of the Gal4-DBD (Suppl. Table 2; FLAG backbone: zfh1-DSCP-3xFLAG-Gal4-dpse-EBFP2). To introduce “3xFLAG-linker”, we performed a mutagenesis PCR using the primers [Phos]ATCGATTACAAGGATGACGATGACAAGGGTGGTGGTGGTAGTATGAAGCTACTGTCTTCTATCGAA and [Phos]GTCATGATCTTTATAATCACCGTCATGGTCTTTGTAGTCCATTTTGAAGTGGCCTGAAGTAAAGGA and the validation plasmid as template (25 μl KAPA HiFi HotStart ReadyMix (KAPA Biosystems KK2602), 1 μl 100 μM forward primer, 1 μl 100 μM reverse primer, template (10 ng/μl), 22 μl double-deionized water; PCR conditions: 95°C 3 min, followed by 21 cycles, 98°C 20 s, 65°C 15 s, 72°C 6 min and final extension 7 min).

After the PCR, the template plasmid was digested using DpnI (Thermo), followed by ligation of the overhanging ends and transformation into Mach1 (Thermo) bacterial cells.

To generate RD expression plasmids with the validation or FLAG backbone, RD fragments amplified from *Drosophila* embryonic cDNA were integrated between SgrDI (Thermo) and BsHTI (Thermo) restriction sites in the respective backbone plasmid via Gibson assembly (NEB) according to the manufacturer’s protocol. “Gal4-DBD control” constructs without an RD were created by annealing the two oligonucleotides CCGGCTGAAGTTGAG and TCGACTCAACTTCAG, encoding two stop codons, and inserting the resulting fragment in between the SgrDI and BsHTI restriction sites of the plasmid backbone.

### *Drosophila* S2 cell culture and cell line generation

#### *Drosophil*a S2 cells were cultured as described before (Arnold *et al*, 2013)

To generate *Drosophila* S2 reporter cell lines, we integrated reporter constructs with 100 bp upstream and downstream homology arms into the integration site at chr2L:9,094,918 which does not contain any genes, by CRISPR-Cas9. The reporter constructs contained 14 UAS sites for Gal4-DBD binding (source: Addgene 128010), an enhancer and a core promoter, the EGFP gene and the SV40 poly-A site. We created 2 different reporters in which EGFP was driven by 1) the zfh1 enhancer and the *Drosophila* synthetic core-promoter (zfh1-DSCP), 2) the ent1 enhancer and the rps12 core-promoter (ent1-rps12) (Suppl. Table 2). These enhancer-promoter pairs were selected based on previous work (Zabidi *et al*, 2015; Arnold *et al*, 2017).

Two plasmids (based on the gRNA expression plasmid Addgene #49330) encoding Cas9 as well as guide RNAs targeting the integration site were kindly received from the Brennecke Lab at IMBA Vienna (Batki *et al*, 2019, chr2L:9,094,918_gRNA_1 and gRNA_2). For the CRISPR-Cas9-mediated integration of the reporters, 50*10^6 *Drosophila* S2 cells were co-transfected with 3.5 μg reporter plasmid and 2.5 μg of each gRNA plasmid using the MaxCyte STX Scalable Transfection System. Cells were passaged for 7 days before selection of GFP-positive cells via fluorescent-activated cell sorting (FACS) and plating in single cell dilutions for generating clonal cell lines. Cells were genotyped using primers binding upstream and downstream of the integration site (Suppl. Table 10: Chr2L_fw and _rv) and homozygous clones were selected.

### Candidate tiling library design and cloning

Candidates for the tiling library were selected based on FlyTF, a database for known and putative *Drosophila melanogaster* transcription factors (Pfreundt *et al*, 2010) in which TFs are scored based on the presence of a DNA-binding domain and experimental evidence for a function in transcription (score of 1-8, with score 1 for the most confident candidates). Of all 1168 FlyTF proteins (for list refer to: http://flytf.gen.cam.ac.uk/flytfmine/begin.do) 1133 factors were selected and 150 bp oligonucleotides were designed to tile the transcripts of these proteins (sliding windows of 6 nt for genes with FlyTF score of 1-4 and sliding window of 15 nt for genes with score of 5-8). This resulted in 209,495 distinct 150 bp candidate fragments (Suppl. Table 1).

The library was cloned from a pool of 209,495 200 bp oligonucleotides synthetized by Twist Biosciences. Each oligonucleotide contained the 150 bp candidate sequence described above flanked by the 25 bp of the partial Illumina i5 (TCCCTACACGACGCTCTTCCGATCT) and 25 bp of the partial i7 (AGATCGGAAGAGCACACGTCTGAAC) adaptor sequences serving as constant linkers for amplification and cloning. The oligonucleotide pool (diluted to 1 ng/µl) was amplified in 40 PCR reactions (98 °C for 45 s; followed by 14 cycles of 98 °C for 15 s, 65 °C for 30 s, 72 °C for 10 s) using KAPA Hifi Hot Start Ready Mix (KAPA Biosystems KK2602) and primers (fw: TTGAGCATGCACCGGACACTCTTTCCCTACACGACGCTCTTCCGATCT and rev: ATCTATCTACGTCGAGTGACTGGAGTTCAGACGTGTGCTCTTCCGATCT) that extended the i5 and i7 adaptor sequences to the full length and added extra 15 bp to each of the adapters, serving as homology arms for directional cloning of the library into RD-seq plasmid (zfh1-DSCP-Gal4-DBD, Suppl. Table 2) vector using In-Fusion HD (Clontech 639650).

### RD-seq pipeline, RNA processing and Illumina sequencing

*Drosophila* S2 reporter cells, cultured at 70-80% confluence, were transfected with the candidate library using the MaxCyte STX Scalable Transfection System. For one screen seven OC-400 processing assemblies were prepared with 200*10^6 cells each in 400 μl MaxCyte Hyclone buffer mixed 1:1 with S2 culture medium without supplements and with 20 μg of the library. In total, for one screen 1.4*10^9 cells were transfected with 140 ug library. S2 cells were electroporated with the pre-set protocol “Optimization 1”, subsequently mixed with 40 μl DNase I (2000 U/ml) in a T175 cell culture flask, incubated for 30 min at 27°C and resuspended in 30 ml complete S2 cell medium.

Three days after transfection, cells were separated into fractions of GFP-positive and - negative cells via fluorescent-activated cell sorting (FACS) on a BD FACSAria III cell sorter. For each experiment 30*10^6 GFP-positive cells and approximately 8*10^6 GFP-negative were collected.

Total RNA of the different fractions was isolated using the Qiagen RNeasy Mini Prep Kit, followed by Poly-A+ RNA enrichment with Dynabeads Oligo-dT25 (Invitrogen) and a DNA digest with TURBO Dnase (Ambion). After RNA cleanup with AMPure XP DNA beads (Agencourt; ratio sample/beads 1:1.8), reverse transcription was performed with Superscript III (50°C for 60 min, 70°C for 15 min; Invitrogen 18080085) and a primer binding within the poly-A site of candidate mRNAs (reverse_transcription_rv: CTCATCAATGTATCTTATCATGTCTG). Next, RNA was digested with Rnase A (Thermo) for 1 h at 37°C, followed by bead cleanup of the cDNA (ratio sample/beads 1:1.4). All subsequent PCR reactions were prepared using the KAPA HiFi HotStart ReadyMix (KAPA Biosystems KK2602). A second strand PCR was performed with a primer binding upstream of the intron sequence which is part of candidate mRNAs (2^nd^_strand_primer_fw: TTGGTAAAGCCACCATGGAAAAG*G) (98°C for 60 s, 65°C for 30 s, 72°C for 90 s), followed by bead cleanup (ratio sample/beads 1:1.4). In the next step, unique molecular identifiers (UMIs) were introduced to the 3’ ends of DNA fragments in a linear PCR with a primer binding to the Illumina i7 adaptor sequence (UMI_primer_rv: CAAGCAGAAGACGGCATACGAGATNNNNNNNNNNGTGACTGGAGTTCAGACGTGT*G) (98°C for 60 s, 65°C for 30 s, 72°C for 90 s). After bead cleanup (ratio sample/beads 1:1.4), the generated fragments were PCR-amplified (98°C 45 s, followed by 16 cycles, 98°C 15 s, 65°C 30 s, 72°C 70s) using two candidate-specific primers (junction_PCR_fw: AAGCCACCATGGAAAAG*G*C*C*A*T and junction_PCR_rv: CAAGCAGAAGACGGCATACG*A), one of which spans the splice junction of the mhc16 intron (5 and 1 nucleotides at the 3’ ends are protected by phosphorothioate bonds, respectively). After another bead cleanup (ratio sample/beads 1:1), candidate fragments were amplified (98°C 45 s, followed by 6-15 cycles, 98°C 15 s, 65°C 30 s, 72°C 70s) with the following primers: i5: aatgatacggcgaccaccgagatctacacXXXXXXXXacactctttccctacacgacgctcttccgatct (XXXXXXXX indicates the position of the index sequence for NGS; for i5 primers used in individual screens, see Suppl. Table 10) and the reverse primer seq_ready_rv: CAAGCAGAAGACGGCATACGAGA*T. PCR products were purified by Agencourt AMPure XP DNA beads (ratio sample/beads 1:0.9), pooled, and subjected to NGS.

All samples were paired-end sequenced (PE36) by the NGS unit of the Vienna Biocenter Core Facilities GmbH (VBCF) on an Illumina NextSeq550 system, following the manufacturer’s protocol.

### Computational analysis of RD-seq hits

#### Creation of dedicated bowtie index

A bowtie index was generated from the designed 150 bp (50 amino acids) oligo sequences, flanked by upstream (“TCCCTACACGACGCTCTTCCGATCT”) and downstream (“AGATCGGAAGAGCACACGTCTGAAC”) adapters. This genome was used to create a custom bowtie index using Bowtie v.1.2.2 (Langmead *et al*, 2009). For visualization purposes in UCSC Genome Browser we also created a linear genome containing selected ordered TFs, separated by 2100 N’s.

#### NGS read mapping and processing

Paired-end sequencing reads were demultiplexed using specific barcodes and mapped to the dedicated bowtie index using Bowtie v.1.2.2 (Langmead *et al*, 2009) (-X 150 ‐v 3 -m 1 – quiet – best – strata). The UMI sequence was incorporated to the read ID at the demultiplexing step. Mapped read pairs, fragments, were collapsed by oligoID and by UMI, i.e., by removing duplicate fragments with identical coordinates if their UMIs differed by <= 2 out of the 10 nucleotides. To calculate position-specific coverage for each frame, oligonucleotide-centric coordinates were transformed into TFs-centric coordinates and total coverage was calculated using the coverage function from R package GenomicRanges v.1.32.7 (Lawrence *et al*, 2013). Fragment coverage was visualized using the linear genome in the UCSC Genome Browser (Kent *et al*, 2002).

We calculated enrichments, hypergeometric P-values, and Benjamini–Hochberg (BH)‐ corrected false discovery rates [FDRs; all statistical calculations done in R (Team RDC, 2008)] between the coverage values in GFP- and GFP+ cells. To define repressive domain (RD) regions, we only considered regions with a minimal coverage of at least 10 fragments in GFP+ and GFP-cells, and selected regions with a minimal enrichment of ≥ 1.5-fold and a hypergeometric P-value of ≤ 1 × 10^−5^ across a minimal length of ≥ 60 bp (20 amino acids), which we extended to include flanking coding sequences (CDS) until P > 1 × 10^−3^ over ≥ 60 bp (20 amino acids).

#### Intersection of RD-seq hits

For each reporter cell line (zfh1-DSP and ent1-rps12) two replicate RD-seq screens were performed. NGS mapping statistics for each screen can be found in (Suppl. Table 11). After determining RD regions for each RD-seq screen, the hits of two replicates were intersected and only repressive regions detected in both replicates with a minimum overlap of 50% were kept for further analysis. Next, repressive regions from the screens with the zfh1-DSCP and the ent1-rps12 reporter cell line were intersected (RD regions with sequence overlaps of 50%, keeping only the longest RD) resulting in 195 unique RD regions which were either detected using both reporter cell lines or only in one of the two. We re-calculated the enrichments of each RD region in each screen to compare their strength between screens and reporters. Information on all 195 RDs can be found in Suppl. Table 3.

### Assessment of sensitivity and specificity of RD-seq

The lack of systematic annotations of RDs in fly TFs makes it difficult to evaluate the specificity and sensitivity of RD-seq against an independent benchmark dataset. However, the candidate library contained fragments covering 438 TFs whose regulatory activity was assessed in a previous study (Stampfel *et al*, 2015). While many TFs could function as repressors in one of the 24 contexts tested by Stampfel *et al*., we defined as repressors TFs that were consistently repressive (sum of scores across all contexts < -20) or strongly repressive (< -35), leading to 156 or 43 TFs, respectively. To allow the assessment of specificity, we additionally defined weakly repressive TFs and non-activators as TFs with sum of scores of < -10 and ≤ 0, respectively. To allow the assessment of sensitivity, we additionally called RDs with a more lenient cutoff of (hypergeometric P-value ≤ 1 × 10−3, minimal enrichment ≥ 1.2-fold). The TFs from Stampfel *et al*. (2015) and RDs detected within these TFs in RD-seq with different cutoffs can be found in Suppl. Table 17.

### RD validations

To validate RD-seq hits, we cloned one of the most highly enriched 150-bp candidates per RD region (sequences in Suppl. Table 4) into the Gal4-DBD validation plasmid backbone zfh1-DSCP-Gal4-dpse-EBFP2 (described in *RD candidate expression plasmids*). All Gibson overhang primers used for the individual RDs can be found in Suppl. Table 10. 25*10^6 reporter cells in 50 μl MaxCyte Hyclone buffer mixed 1:1 with S2 culture medium without supplements were transfected with 2.5 μg Gal4-DBD-RD or Gal4-DBD control plasmid using OC-100 processing assemblies and the MaxCyte STX Scalable Transfection System on “Optimization 1”. After electroporation, cells were resuspended in 5 μl DNase I (2000 U/ml) in a T25 cell culture flask, incubated for 30 min at 27°C and resuspended in 5 ml complete S2 cell medium.

Three days after transfection cells were submitted to flow cytometry analysis using a FACS BD LSR Fortessa (BD Biosciences). The GFP signal of transfected cells, gated based on EBFP2 expression as transfection control, was determined and data analysis was performed with FACS Diva. As a measure of the repressive strength of the RD, we used the ratio of the medians between the GFP signal of cells expressing a Gal4-DBD control construct without a RD and cells expressing the Gal4-DBD-RD and called it fold change (FC) repression (FC repression = median-GFP[Gal4-DBD control]/ median-GFP[Gal4-DBD-RD]). We used two-tailed Student T-tests to assess the significance of the difference to Gal4-DBD for each RD (P<=0.05; FC>1 for validated). FC repression values from individual replicates and P-values of the T-tests can be found in Suppl. Table 4.

### Analysis of RD and DBD positioning within full-length proteins

We used the centered amino acid of each 50 AA RD (RD-seq) and DBD (from ProSitePatterns and Pfam) as their position within the full-length TF sequences, scaled over the length of the respective protein sequences to be comparable across proteins. To analyze DBD positioning we only considered DBDs appearing in proteins that have an RD region according to RD-seq (Suppl. Table 12).

### Analysis of RD overlaps with known domains and IDRs

We used the full-length protein sequences of all proteins for which an RD was detected in RD-seq as input for ProSitePatterns (de Castro *et al*, 2006), Pfam (Mistry *et al*, 2021) and MobiDB-lite (Necci *et al*, 2021) protein domain database searches. To assign a ProSitePatterns, Pfam or MobiDB-lite hit to a RD, we only selected those cases in which the RD (=50 AA most strongly enriched candidate fragment within the RD region) contains at least 50% of the domain or in which at least 50% of the RD is part of the annotated domain. ProSitePatterns and Pfam entries from protein families, not relevant for protein domain analysis, were removed. The resulting domain–RD overlaps can be found in Suppl. Table 5.

### MEME and FIMO peptide motif searches among RD-seq hits

The most repressive 150 bp candidate fragments (= 50 AA long RDs) were used for MEME *de novo* motif analyses. For that, 4 different sets of RDs were created based on the preference of an RD region for the zfh1-DSCP or the ent1-rps12 reporter context. Preferences for one of the reporters were calculated by dividing the mean FC of the RD region detected in the RD-seq screens using one reporter over the FC resulting from the RD-seq screens with the other reporter. Subset information can be found in Suppl. Table 3 in the column “RD.region.preference.1.3fold”. RD regions with a >1.3-fold preference for the zfh1-DSCP context were categorized as “zfh1” hits, while RD regions with a >1.3-fold preference for the ent1-rps12 reporter were categorized as “ent1” hits. RDs without a preference were categorized as “global” hits. This resulted in 4 different RD sets that were separately subjected to MEME *de novo* motif searches (Bailey *et al*, 2015): i) 195 RDs (all hits), ii) 89 RDs without a preference, iii) 43 zfh1 RDs, iv) 63 ent1 RDs.

We ran MEME v.5.1.1 (Bailey *et al*, 2015) with the following parameters: *-protein -oc*. *-nostatus -time 18000 -mod zoops -nmotifs 25 -minw 4 -maxw 15 -objfun classic - markov_order 0*. This resulted in 22 motifs in each set with motif widths between 4 and 15 AA. Two motifs were removed since the enrichment derived solely from paralog proteins. To collapse redundant motifs by similarity, we computed the distances between all motif pairs using TOMTOM (kullback distance) (Gupta *et al*, 2007) and performed hierarchical clustering using Pearson correlation as the distance metric and complete linkage using the hclust R function. The tree was cut at height 0.7, resulting in 11 non-redundant motif clusters that were manually annotated (Fig. 2C and Suppl. Table 6). Some of the motifs were detected in multiple RD sets (e.g. EH1 motif was found in MEME searches with zfh1, global and all RDs, see Fig. 2C). Hence, for subsequent analysis we selected one motif per group: Motif 1 – ent1, Motif 2 – all, Motif 3 – global, Motif 4 – global, Motif 5 – global, Motif 6 – zfh1, Motif 7 – all, Motif 8 – zfh1, Motif 9 – all, Motif 10 – all, Motif 11 – ent1. These MEME motifs were used as input for FIMO searches (v.5.4.1.) (Grant *et al*, 2011) with a stringent (p<0.0001) or a lenient (p<0.001) cutoff to determine the prevalence of the peptide motifs among all 195 RD-seq hits. The results of the FIMO searches can be found in Suppl. Table 6.

### Analysis of known SLiMs within RDs using the ELM prediction tool

We used the most repressive 150 bp candidate fragment (= 50 AA long RDs) within each of the 195 RD regions detected in RD-seq as input for ELM database searches for short linear motifs (SLiMs) (Kumar *et al*, 2019). Next, we used the list of matches to high-probability ELM patterns (p<0.0002) and filtered for SLiMs that have been implicated in the interaction with co-repressors. These were the EH1 motif (LIG_EH1_1), the WRPW motif (LIG_WRPW_2), the CtBP ligand motif (LIG_CtBP_PxDLS_1), the Sin3A-interacting domain (LIG_Sin3_1) and the HCF-1 binding motif (LIG_HCF-1_HBM_1) (Suppl. Table 7).

### Analysis of known and novel SLiMs within RDs

We characterized the motif composition of each RD by integrating both annotated (from ELM) and *de novo* (from MEME) SLiMs (Fig. 2 D). We categorized an RD as having a known SLiM instance if containing an instance from ELM, while the remaining RDs with instances from MEME analysis not reported in ELM were considered as novel instances. The remaining RDs without any of these SLiMs were considered as unexplained.

### Site-directed mutagenesis of RD peptide motifs

To determine the requirement of peptide motifs discovered in MEME and FIMO searches for the function of RDs, residues within these motifs were mutated to Alanines (5 AA mutated to Ala in case of EH1, PXDLS, AAxxL and PLKKR motifs, and 4 AA in case of the HKKF motif). The Gal4-DBD-RD validation plasmids with the wild type RD sequences were subjected to site-directed mutagenesis using primers carrying the mutated version of the motifs in overhangs (primers see Suppl. Table 10). After PCR amplification with the KAPA HiFi HotStart ReadyMix (KAPA Biosystems KK2602) (95°C 3 min, followed by 21 cycles, 98°C 20 s, 65°C 15 s, 72°C 6 min and final extension 7 min), amplicons were purified using the NEB Monarch Gel Extraction kit and template plasmids were DpnI-digested (Thermo) followed by cleanup with the NEB Monarch Nucleic Acid kit. The ends created by the overhang primers were ligated and Mach1 cells (Thermo) were transformed with the resulting plasmids. Mutated Gal4-DBD-RD constructs were used in validation experiments as described above. Wild type and mutant RD sequences and the validation results can be found in Suppl. Table 4.

### Assessing RD expression in western blots

To monitor the expression of mutated RDs in comparison to the wild type RDs, wild type and mutant RDs were cloned into the FLAG-Gal4-DBD background (Suppl. Table 2, zfh1-DSCP-3xFLAG-Gal4-dpse-EBFP2) as described under *RD candidate expression plasmids*. The zfh1-DSCP reporter cell line was transfected with the FLAG-Gal4-DBD-RD plasmids according to *RD validations*. Three days after transfection, 3*10^6 cells were harvested, washed with PBS, and lysed in 30 μl lysis buffer (10 mM Tris pH8, 1 mM EDTA, 0.5 mM EGTA, 1% Triton x-100, 0.1% SDS, 0.1% sodium deoxycholate, 140 mM NaCl, Roche cOmplete Protease Inhibitor, Benzonase (Sigma, 2.5 Units/μl)) for 10 min on ice. 30 μl 2x Laemmli Sample Buffer (Bio-Rad) with 5% b-mercaptoethanol were added to the sample followed by incubation at 95°C for 5 min. Proteins were separated using SDS-polyacrylamide gel electrophoresis (Bio-Rad) and subsequently blotted onto a 0.2 μm nitrocellulose membrane (Power Blotter XL, Invitrogen). The membrane was blocked with 5% milk in TBS-T (TBS with 1% Tween-20) and incubated over night at 4°C with the primary anti-FLAG antibody (Sigma F1804-200UG, 1:1000 in 2.5% milk in TBS-T). The membrane was washed 3 times with TBS-T, followed by 1 h incubation with the HRP-conjugated secondary antibody (Cell Signaling 7076S, 1:10,000 in 2.5% milk in TBS-T). After three washes in TBS-T, the membrane was incubated with Clarity Western ECL Blotting Substrate (Bio-Rad) and imaged with a ChemiDoc MP imaging system (Bio-Rad). For a loading control, blots were probed with a primary anti-Tubulin antibody (Abcam, ab18251).

### Immunoprecipitation-Mass Spectrometry (IP-MS) experiments

RDs were cloned into the zfh1-DSCP-3xFLAG-Gal4-dpse-EBFP2 plasmid backbone as described under *RD candidate expression plasmids*. Plasmids encoding RDs with a specific peptide motif were mixed in an equal molar ratio to create RD plasmid pools (PxDLS: CG42741-RD, Tio-RD, Ham-RD, CG11122-RD1; AAxxL: CG11617-RD2, Cic-RD2, Glut4EF-RD, CG12605-RD; PLKKR: Ash1-RD, Kr-h1-RD2, Net-RD, Vri-RD; HKKF: Eip75B-RD, CHES-1-like-RD1, Kah-RD, Shn-RD1). As a control we used a 3xFLAG-Gal4-DBD construct without an RD sequence. For each replicate of an IP-MS experiment 200*10^6 *Drosophila* S2 cells in 400 μl MaxCyte Hyclone buffer mixed 1:1 with S2 culture medium without supplements were transfected with 30 μg 3xFLAG-Gal4-DBD control plasmid or 30 μg of an RD plasmid pool using OC-400 processing assemblies and the MaxCyte STX Scalable Transfection System on “Optimization 1”. After electroporation, cells were resuspended in 40 μl DNase I (2000 U/ml) in a T175 cell culture flask, incubated for 30 min at 27°C and resuspended in 30 ml complete S2 cell medium.

One day after transfection, cells were harvested, washed in PBS and incubated in buffer A (10 mM Tris pH 7.5, 2 mM MgCl_2_, 3 mM CaCl2, Sigma cOmplete EDTA-free Protease Inhibitor Cocktail) for 15 min at 4 °C followed by centrifugation. The pellet was resuspended and incubated for 30 min at 4 °C in buffer B (10 mM Tris pH 7.5, 2 mM MgCl_2_, 3 mM CaCl2, 0.5% IGEPAL CA-630, 10% Glycerol, 1 mM DTT, Sigma cOmplete EDTA-free Protease Inhibitor Cocktail). After centrifugation, the nuclear pellet was resuspended in buffer C (40 mM HEPES pH 7.6, 4 mM MgCl_2_, 0.6% Triton X-100, 0.5% IGEPAL CA-630, 20% Glycerol, 1 mM DTT, Sigma cOmplete EDTA-free Protease Inhibitor Cocktail) with 100 mM NaCl and incubated for 30 min at 4 °C, followed by centrifugation. The supernatant containing the nucleoplasm was collected and the remaining chromatin pellet was resuspended in buffer C with 300 mM NaCl and subjected to sonication with a Diagenode Bioruptor Sonicator for 10 min at low intensity. After centrifugation the supernatant was transferred to the nucleoplasmic fraction. FLAG M2 Magnetic Beads (Sigma, M8823) were equilibrated in buffer C with 150 mM NaCl. Nuclear lysate was added to the beads for immunoprecipitation over night at 4 °C. Afterwards, the beads were washed 3 times in buffer C with 150 mM NaCl, followed by 4 washes in non-detergent buffer (20 mM Tris pH 7.5, 130 mM NaCl).

Beads were resuspended in 80 μl of 100 mM ammonium bicarbonate (ABC), supplemented with 800 ng of lysyl endopeptidase (Lys-C, Fujifilm Wako Pure Chemical Corporation) and incubated for 4 hours on a Thermo-shaker with 1200 rpm at 37°C. The supernatant was transferred to a fresh tube and reduced with 1 mM Tris 2-carboxyethyl phosphine hydrochloride (TCEP, Sigma) for 30 minutes at 60°C and alkylated in 4 mM methyl methanethiosulfonate (MMTS, Fluka) for 30 min at room temperature. Subsequently, the sample was digested with 800 ng trypsin (Trypsin Gold, Promega) at 37°C over night. The digest was acidified by addition of trifluoroacetic acid (TFA, Pierce) to 1%. A similar aliquot of each sample was analysed by LC-MS/MS.

#### nanoLC-MS/MS Analysis

The nano HPLC system (UltiMate 3000 RSLC nano system, Thermo Fisher Scientific) was coupled to an Exploris 480 mass spectrometer equipped with a FAIMS pro interfaces and a Nanospray Flex ion source (all parts Thermo Fisher Scientific). Peptides were loaded onto a trap column (PepMap Acclaim C18, 5 mm × 300 μm ID, 5 μm particles, 100 Åpore size, Thermo Fisher Scientific) at a flow rate of 25 μl/min using 0.1% TFA as mobile phase. After 10 minutes, the trap column was switched in line with the analytical column (PepMap Acclaim C18, 500 mm × 75 μm ID, 2 μm, 100 Å, Thermo Fisher Scientific) operated at 30°C. Peptides were eluted using a flow rate of 230 nl/min, starting with the mobile phases 98% A (0.1% formic acid in water) and 2% B (80% acetonitrile, 0.1% formic acid) and linearly increasing to 35% B over the next 120 minutes.

The Exploris mass spectrometer was operated in data-dependent mode, performing a full scan (m/z range 350-1200, resolution 60,000, target value 1E6) at 3 different compensation voltages (CV-45, -60, -75), followed each by MS/MS scans of the most abundant ions for a cycle time of 0.9 (CV -45, -60) or 0.7 (CV -75) seconds per CV. MS/MS spectra were acquired using a collision energy of 30, isolation width of 1.0 m/z, resolution of 30.000, target value of 2E5 and intensity threshold of 2.5E4, maximum injection time 100 ms. Precursor ions selected for fragmentation (include charge state 2-6) were excluded for 45 s. The monoisotopic precursor selection filter and exclude isotopes feature were enabled.

#### IP-MS data processing

For peptide identification, the RAW-files were loaded into Proteome Discoverer (version 2.5.0.400, Thermo Scientific). All MS/MS spectra were searched using MSAmanda v2.0.0.16129 (Dorfer *et al*, 2014). The peptide and fragment mass tolerance was set to ±10 ppm, the maximal number of missed cleavages was set to 2, using tryptic enzymatic specificity without proline restriction. Peptide and protein identification was performed in two steps. For an initial search the RAW-files were searched against the database dmel-all-translation-r6.43.fasta (Flybase.org, 22,232 sequences; 20,321,723 residues), supplemented with common contaminants and sequences of tagged proteins of interest, using the following search parameters: beta-methylthiolation of cysteine was set as a fixed modification, oxidation of methionine as variable modification. The result was filtered to 1 % FDR on protein using the Percolator algorithm (Käll *et al*, 2007) integrated in Proteome Discoverer. A sub-database of proteins identified in this search was generated for further processing. For the second search, the RAW-files were searched against the created sub-database using the same settings as above plus considering additional variable modifications: Phosphorylation on serine, threonine and tyrosine, deamidation on asparagine and glutamine, and glutamine to pyro-glutamate conversion at peptide N-terminal glutamine, acetylation on protein N-terminus were set as variable modifications. The localization of the post-translational modification sites within the peptides was performed with the tool ptmRS, based on the tool phosphoRS (Taus *et al*, 2011). Identifications were filtered again to 1 % FDR on protein and PSM level, additionally an Amanda score cut-off of at least 150 was applied. Peptides were subjected to label-free quantification using IMP-apQuant (Doblmann *et al*, 2018). Proteins were quantified by summing unique and razor peptides or only unique peptides and applying intensity-based absolute quantification (iBAQ) (Schwanhäusser *et al*, 2011). FLAG-Gal4-DBD-RD bait proteins were filtered to be identified by a minimum of 2 PSMs in at least 1 sample. All other proteins were filtered to be identified by a minimum of 3 quantified peptides in at least 1 sample. Protein-abundances-normalization was done using sum normalization. Differential abundance protein analysis between each RD group and Gal4-DBD constructs was performed using limma (Smyth, 2004), considering all replicates. The results of the differential abundance analysis can be found in Suppl. Table 13.

### RNAi-mediated depletion of co-repressors

For RNAi-mediated depletion of CoRs, two distinct long dsRNAs targeting each CoR, without off-target effects were selected from UP-TORR (Hu *et al*, 2013) (https://www.flyrnai.org/up-torr/). As a negative control we used a dsRNA targeting the Renilla Luciferase which is not expressed in *Drosophila* S2 cells (sequences in Suppl. Table 14). Primers including the T7 promoter sequence (TAATACGACTCACTATAGGG) in their overhangs (Suppl. Table 14) were used to amplify these dsRNA-complementary sequences from *Drosophila* genomic DNA with the Q5® Hot Start High-Fidelity 2X Master Mix (NEB). The PCR product was precipitated in 1 volume isopropanol and 1/10 3M sodium acetate for 5 min at room temperature, followed by centrifugation for 20 min at 18,000 g at 4°C, a wash with 70% ethanol and resuspension in nuclease-free water. Subsequently, the fragments were transcribed with the T7 RNA Polymerase (Promega) at 37°C over night. After DNase digest (Turbo DNase I Ambion) at 37°C for 1 h the RNA was purified in a phenol-chloroform extraction. Samples were treated with 1 volume of Acid-Phenol-Chloroform (Roti-Aqua-P/C/I) for 5 min at room temperature followed by centrifugation and recovery of the aqueous phase. The RNA was precipitated by adding 2.5 volumes 100% ethanol and 1/10 volume 3 M sodium acetate, incubation at -20°C for 30 min. After centrifugation and washing of with 70% ethanol, the RNA was purified using the Invitrogen MEGAclear Transcription Clean-Up Kit.

*Drosophila* zfh1-DSCP reporter cells were transfected with the Gal4-DBD-RD plasmid or the Gal4-DBD control plasmid according to *RD validations*, but using 5 μg instead of 2.5 μg plasmid for 25*10^6 cells. 16 h after transfection, cells were harvested, washed twice in PBS and resuspended in serum-free medium (ExpressFive SFM (Invitrogen), 16 mM Glutamine (Gibco)). For each condition 0.75*10^6 cells in 500 μl serum-free medium were seeded into 12-well tissue culture plates, 20 μg dsRNA were added and incubated for 1 h at 27°C, before adding 1 ml full medium (ExpressFive SFM Invitrogen, 16 mM Glutamine, 10% FBS (Sigma-Aldrich), 1% penicillin-streptomycin (Gibco)) to each well. Three days after dsRNA treatment, cells were submitted to flow cytometry analysis as in *RD validations*. The fold-change (FC) repression was determined as the ratio of the median GFP signal of transfected cells compared between cells expressing the Gal4-DBD control and cells expressing the Gal4-DBD-RD, both treated with the same dsRNA. FC repression values and results of two-tailed Student T-tests can be found in Suppl. Table 14.

Reverse transcription quantitative PCR (RT-qPCR) was performed to assess the depletion of the endogenous co-repressors. Three days after treatment of non-transfected reporter cells as described above, cells were harvested, followed by total RNA isolation with the Quiagen RNeasy Mini Kit and DNA digest with Ambion Turbo DNaseI. The RNA was reverse transcribed using Oligo(dt)20 primer (Invitrogen, 18418020) and SuperScript III Reverse Transcriptase (Invitrogen). qPCR with three technical replicates per condition was performed with the Promega GoTaq qPCR Master Mix (qPCR primers in Suppl. Table 14). qPCR was analyzed using the Delta-Delta Ct Method (Livak & Schmittgen, 2001). Conditions with primers targeting the rps12 gene were used as a housekeeping gene control. In brief, the following equations were used: DeltaCt = mean Ct CoR primers – mean Ct rps12 primers; DeltaDeltaCt = DeltaCt – RenillaDeltaCt; FC = 2^(-DeltaDelatCt).

### Sequence alignments for RD-containing repressors

Orthologs of *Drosophila* proteins harboring RDs with specific repressive motifs were detected in the NCBI protein or UniProt reference database, based on NCBI blast searches applying significant e-values (< 0.001) and considering reciprocal best hits (Agarwala *et al*, 2018; Bateman *et al*, 2021; Altschul, 1997). In addition to fruit fly (*Drosophila melanogaster*), 6 other species were selected for a long evolutionary distance and presence in all 4 motifs, namely southern house mosquito (*Culex quinquefasciatus*), American lobster (*Homarus americanus*), a tardigrade (*Ramazzottius varieornatus*), a bivalve (*Mytilus coruscus*), zebrafish (*Danio rerio*), and human (*Homo sapiens*). Alignments were performed with mafft (-linsi, v7.427) (Katoh & Toh, 2008) and visualization in Jalview (ClustalX coloring scheme) (Waterhouse *et al*, 2009). Accessions and gene names are given in Suppl. Table 15. Gene names are according to Uniprot or NCBI nomenclature.

### Analysis of motif conservation in fly and human proteins

To measure the conservation of each amino acid of *Drosophila melanogaster* and human transcription-related proteins, we first identified groups of orthologous proteins (= orthogroups) across a range of species from either the Panarthropoda clade for comparison to *Drosophila*, or the vertebrate clade for comparison to human with Orthofinder (Emms & Kelly, 2019) and used these groups for multiple sequence alignments.

64 species of the Panarthropoda clade, and 40 species from the vertebrate clade were selected from the UniProt reference proteomes (Bateman *et al*, 2021) (Suppl. Table 16). Orthogroups were detected using OrthoFinder for the clades individually, with diamond ultra-sensitive mode and an e-value threshold of 0.001, version 2.5.4 (Emms & Kelly, 2019).

In the Panarthropoda set, 590 orthogroups had all species present, and were used to infer a rooted species tree with STAG and to build hierachical orthogroups (HOGs) in OrthoFinder (preprint: Emms and Kelly, 2018). We used the list of 1133 transcription-related proteins from *Drosophila melanogaster* (Suppl. Table 1). We only processed orthogroups containing equal or less than 150 entries and 1024 orthogroups of the root node (N0, Panarthropoda). 1072 of the *Drosophila* transcription-related proteins, fulfilled these criteria. Four more orthogroups (9 transcription factors) were derived from the N2 node (insects), and one more orthogroup from the N6 node (Endopterygota).

In the vertebrates set, 3775 orthogroups contained all species and were used for the species tree. The human transcription factor list contained 2754 IDs (Suppl. Table 8) that were mapped to 2740 UniProt entries. 2259 orthogroups (2470 UniProt IDs) were retrieved from the root N0 (vertebrates) node, 29 orthogroups (116 IDs) with the N6 node (tetrapods), and 5 orthogroups (38 IDs) with the N14 node (mammals).

All orthogroup sequences were aligned with mafft (-linsi mode, v7.427) (Katoh & Toh, 2008) and the sequence conservation score calculated with AAcon (KARLIN method, results normalized with values between 0 and 1) (see Golicz *et al*, 2018, AACon: A Fast Amino Acid Conservation Calculation Service. Submitted paper. http://www.compbio.dundee.ac.uk/aacon/).

We next mapped the positions of all instances of the five main SLiMs (EH1, PLKKR, HKKF, PxDLS, AAxxL) within the protein sequence of *Drosophila* and human transcription-related factors using FIMO (as described in section *MEME and FIMO peptide motif searches among RD-seq hits*). We quantified the conservation of each instance as the averaged conservation of its amino acids and compared it with the average conservation of the flanking amino acids (sequences with same total length as the motifs up- and downstream of motif instance) (Fig. 4 C, D).

### FIMO searches among human transcription-related proteins

In order to predict RDs in human proteins we used the PxDLS, PLKKR, EH1, HKKF, AAxxL motifs (same motifs as used for FIMO searches among 195 RDs, see section *MEME and FIMO peptide motif searches among RD-seq hits*) found in fly as input for FIMO searches (v.5.4.1) (Grant *et al*, 2011) among human transcription-related genes (based on Lambert *et al*, 2018; Vaquerizas *et al*, 2009) (Suppl. Table 8). The results of the FIMO searches for the PxDLS and PLKKR motifs among human transcription-related genes can be found in Suppl. Table 9.

## Data availability

Raw sequencing data will be made available on online repositories. Mass spectrometry raw data as well as *Drosophila* and human protein conservation scores can be found on zenodo at https://doi.org/10.5281/zenodo.6786955. Genome browser tracks showing all read coverage tracks and RD regions for the different screens are available at https://genome.ucsc.edu/s/bernardo.almeida/RDseq_manuscript.

## References

Agarwala R, Barrett T, Beck J, Benson DA, Bollin C, Bolton E, Bourexis D, Brister JR, Bryant SH, Canese K, et al (2018) Database resources of the National Center for Biotechnology Information. Nucleic Acids Res 46: D8–D13

Alerasool N, Leng H, Lin ZY, Gingras AC & Taipale M (2022) Identification and functional characterization of transcriptional activators in human cells. Mol Cell 82: 677-695.e7

Alerasool N, Segal D, Lee H & Taipale M (2020) An efficient KRAB domain for CRISPRi applications in human cells. Nat Methods 17: 1093–1096

Altschul S (1997) Gapped BLAST and PSI-BLAST: a new generation of protein database search programs. Nucleic Acids Res 25: 3389–3402

Arnold CD, Gerlach D, Stelzer C, Boryń ŁM, Rath M & Stark A (2013) Genome-wide quantitative enhancer activity maps identified by STARR-seq. Science (80-) 339: 1074–1077

Arnold CD, Nemčko F, Woodfin AR, Wienerroither S, Vlasova A, Schleiffer A, Pagani M, Rath M & Stark A (2018) A high-throughput method to identify trans-activation domains within transcription factor sequences. EMBO J: e98896

Arnold CD, Zabidi MA, Pagani M, Rath M, Schernhuber K, Kazmar T & Stark A (2017) Genome-wide assessment of sequence-intrinsic enhancer responsiveness at single-base-pair resolution. Nat Biotechnol 35: 136–144

Atanesyan L, Günther V, Dichtl B, Georgiev O & Schaffner W (2012) Polyglutamine tracts as modulators of transcriptional activation from yeast to mammals. Biol Chem 393: 63–70

Bailey TL, Johnson J, Grant CE & Noble WS (2015) The MEME Suite. Nucleic Acids Res 43: W39–W49

Basu S, Mackowiak SD, Niskanen H, Knezevic D, Asimi V, Grosswendt S, Geertsema H, Ali S, Jerković I, Ewers H, et al (2020) Unblending of Transcriptional Condensates in Human Repeat Expansion Disease. Cell 181: 1062-1079.e30

Bateman A, Martin M-J, Orchard S, Magrane M, Agivetova R, Ahmad S, Alpi E, Bowler-Barnett EH, Britto R, Bursteinas B, et al (2021) UniProt: the universal protein knowledgebase in 2021. Nucleic Acids Res 49: D480–D489

Batki J, Schnabl J, Wang J, Handler D, Andreev VI, Stieger CE, Novatchkova M, Lampersberger L, Kauneckaite K, Xie W, et al (2019) The nascent RNA binding complex SFiNX licenses piRNA-guided heterochromatin formation. Nat Struct Mol Biol 26: 720–731

Belacortu Y, Weiss R, Kadener S & Paricio N (2012) Transcriptional activity and nuclear localization of cabut, the Drosophila ortholog of vertebrate TGF-β-Inducible Early-Response gene (TIEG) proteins. PLoS One 7

Boija A, Klein IA, Sabari BR, Dall’Agnese A, Coffey EL, Zamudio A V., Li CH, Shrinivas K, Manteiga JC, Hannett NM, et al (2018) Transcription Factors Activate Genes through the Phase-Separation Capacity of Their Activation Domains. Cell: 1–14

Brayer KJ & Segal DJ (2008) Keep Your Fingers Off My DNA: Protein–Protein Interactions Mediated by C2H2 Zinc Finger Domains. Cell Biochem Biophys 50: 111–131

Brennecke J, Stark A, Russell RB & Cohen SM (2005) Principles of MicroRNA–Target Recognition. PLoS Biol 3: e85

Brent R & Ptashne M (1985) A eukaryotic transcriptional activator bearing the DNA specificity of a prokaryotic repressor. Cell 43: 729–736

Brodsky S, Jana T, Mittelman K, Chapal M, Kumar DK, Carmi M & Barkai N (2020) Intrinsically Disordered Regions Direct Transcription Factor In Vivo Binding Specificity. Mol Cell 79: 459-471.e4

Cai Y & Laughon A (2009) The Drosophila Smad cofactor Schnurri engages in redundant and synergistic interactions with multiple corepressors. Biochim Biophys Acta - Gene Regul Mech 1789: 232–245

de Castro E, Sigrist CJA, Gattiker A, Bulliard V, Langendijk-Genevaux PS, Gasteiger E, Bairoch A & Hulo N (2006) ScanProsite: detection of PROSITE signature matches and ProRule-associated functional and structural residues in proteins. Nucleic Acids Res 34: W362–W365

Chaubal A & Pile LA (2018) Same agent, different messages: insight into transcriptional regulation by SIN3 isoforms. Epigenetics Chromatin 11: 17

Chinnadurai G (2002) CtBP, an unconventional transcriptional torepressor in development and oncogenesis. Mol Cell 9: 213–224

Chong S, Dugast-Darzacq C, Liu Z, Dong P, Dailey GM, Cattoglio C, Heckert A, Banala S, Lavis L, Darzacq X, et al (2018) Imaging dynamic and selective low-complexity domain interactions that control gene transcription. Science (80-) 361: eaar2555

Copley RR (2005) The EH1 motif in metazoan transcription factors. BMC Genomics 6: 1–8

Doblmann J, Dusberger F, Imre R, Hudecz O, Stanek F, Mechtler K & Dürnberger G (2018) apQuant: Accurate Label-Free Quantification by Quality Filtering. J Proteome Res: acs.jproteome.8b00113

Dorfer V, Pichler P, Stranzl T, Stadlmann J, Taus T, Winkler S & Mechtler K (2014) MS Amanda, a Universal Identification Algorithm Optimized for High Accuracy Tandem Mass Spectra. J Proteome Res 13: 3679–3684

Emms DM & Kelly S (2019) OrthoFinder: phylogenetic orthology inference for comparative genomics. Genome Biol 20: 238

Erijman A, Kozlowski L, Sohrabi-Jahromi S, Fishburn J, Warfield L, Schreiber J, Noble WS, Söding J & Hahn S (2020) A High-Throughput Screen for Transcription Activation Domains Reveals Their Sequence Features and Permits Prediction by Deep Learning. Mol Cell 78: 890-902.e6

Erkina TY & Erkine AM (2016) Nucleosome distortion as a possible mechanism of transcription activation domain function. Epigenetics Chromatin 9: 40

Fisher AL, Ohsako S & Caudy M (1996) The WRPW motif of the hairy-related basic helix-loop-helix repressor proteins acts as a 4-amino-acid transcription repression and protein-protein interaction domain. Mol Cell Biol 16: 2670–2677

Grant CE, Bailey TL & Noble WS (2011) FIMO: scanning for occurrences of a given motif. Bioinformatics 27: 1017–1018

Gupta S, Stamatoyannopoulos JA, Bailey TL & Noble W (2007) Quantifying similarity between motifs. Genome Biol 8: R24

Han K & Manley JL (1993a) Functional domains of the Drosophila Engrailed protein. EMBO J 12: 2723–2733

Han K & Manley JL (1993b) Transcriptional repression by the Drosophila even-skipped protein: Definition of a minimal repression domain. Genes Dev 7: 491–503

Hanna-Rose W, Licht JD & Hansen U (1997) Two evolutionarily conserved repression domains in the Drosophila Kruppel protein differ in activator specificity. Mol Cell Biol 17: 4820–9.

He X, Nie Y, Zhou H, Hu R, Li Y, He T, Zhu J, Yang Y & Liu M (2021) Structural Insight into the Binding of TGIF1 to SIN3A PAH2 Domain through a C-Terminal Amphipathic Helix. Int J Mol Sci 22: 12631

Hemavathy K, Hu X, Ashraf SI, Small SJ & Ip YT (2004) The repressor function of Snail is required for Drosophila gastrulation and is not replaceable by Escargot or Worniu. Dev Biol 269: 411–420

Hu Y, Roesel C, Flockhart I, Perkins L, Perrimon N & Mohr SE (2013) UP-TORR: Online Tool for Accurate and Up-to-Date Annotation of RNAi Reagents. Genetics 195: 37–45

Izutsu K, Kurokawa M, Imai Y, Maki K, Mitani K & Hirai H (2001) The corepressor CtBP interacts with Evi-1 to repress transforming growth factor β signaling. Blood 97: 2815–2822

Jennings BH & Ish-Horowicz D (2008) The Groucho/TLE/Grg family of transcriptional co-repressors. Genome Biol 9: 205

Jennings BH, Pickles LM, Wainwright SM, Roe SM, Pearl LH & Ish-Horowicz D (2006) Molecular Recognition of Transcriptional Repressor Motifs by the WD Domain of the Groucho/TLE Corepressor. Mol Cell 22: 645–655

Kajimura S, Seale P, Tomaru T, Erdjument-Bromage H, Cooper MP, Ruas JL, Chin S, Tempst P, Lazar MA & Spiegelman BM (2008) Regulation of the brown and white fat gene programs through a PRDM16/CtBP transcriptional complex. Genes Dev 22: 1397–1409

Käll L, Canterbury JD, Weston J, Noble WS & MacCoss MJ (2007) Semi-supervised learning for peptide identification from shotgun proteomics datasets. Nat Methods 4: 923–925

Katoh K & Toh H (2008) Recent developments in the MAFFT multiple sequence alignment program. Brief Bioinform 9: 286–298

Katz SG, Cantor AB & Orkin SH (2002) Interaction between FOG-1 and the Corepressor C-Terminal Binding Protein Is Dispensable for Normal Erythropoiesis In Vivo. Mol Cell Biol 22: 3121–3128

Kent WJ, Sugnet CW, Furey TS, Roskin KM, Pringle TH, Zahler AM & Haussler and D (2002) The Human Genome Browser at UCSC. Genome Res 12: 996–1006

Kruusvee V, Lyst MJ, Taylor C, Tarnauskaitė Ž, Bird AP & Cook AG (2017) Structure of the MeCP2–TBLR1 complex reveals a molecular basis for Rett syndrome and related disorders. Proc Natl Acad Sci 114

Kumar M, Gouw M, Michael S, Sámano-Sánchez H, Pancsa R, Glavina J, Diakogianni A, Valverde JA, Bukirova D, Ĉalyševa J, et al (2019) ELM—the eukaryotic linear motif resource in 2020. Nucleic Acids Res

Lambert SA, Jolma A, Campitelli LF, Das PK, Yin Y, Albu M, Chen X, Taipale J, Hughes TR & Weirauch MT (2018) The Human Transcription Factors. Cell 172: 650–665

Langmead B, Trapnell C, Pop M & Salzberg SL (2009) Ultrafast and memory-efficient alignment of short DNA sequences to the human genome. Genome Biol 10: R25

Lawrence M, Huber W, Pagès H, Aboyoun P, Carlson M, Gentleman R, Morgan MT & Carey VJ (2013) Software for Computing and Annotating Genomic Ranges. PLoS Comput Biol 9: e1003118

Lee J-A, Suh D-C, Kang J-E, Kim M-H, Park H, Lee M-N, Kim J-M, Jeon B-N, Roh H-E, Yu M-Y, et al (2005) Transcriptional Activity of Sp1 Is Regulated by Molecular Interactions between the Zinc Finger DNA Binding Domain and the Inhibitory Domain with Corepressors, and This Interaction Is Modulated by MEK. J Biol Chem 280: 28061–28071

Lewis BP, Burge CB & Bartel DP (2005) Conserved Seed Pairing, Often Flanked by Adenosines, Indicates that Thousands of Human Genes are MicroRNA Targets. Cell 120: 15–20

Livak KJ & Schmittgen TD (2001) Analysis of Relative Gene Expression Data Using Real-Time Quantitative PCR and the 2−ΔΔCT Method. Methods 25: 402–408

Logan C, Hanks MC, Noble-Topham S, Nallainathan D, Provart NJ & Joyner AL (1992) Cloning and sequence comparison of the mouse, human, and chicken engrailed genes reveal potential functional domains and regulatory regions. Dev Genet 13: 345–358

Lyst MJ, Ekiert R, Ebert DH, Merusi C, Nowak J, Selfridge J, Guy J, Kastan NR, Robinson ND, de Lima Alves F, et al (2013) Rett syndrome mutations abolish the interaction of MeCP2 with the NCoR/SMRT co-repressor. Nat Neurosci 16: 898–902

Mistry J, Chuguransky S, Williams L, Qureshi M, Salazar GA, Sonnhammer ELL, Tosatto SCE, Paladin L, Raj S, Richardson LJ, et al (2021) Pfam: The protein families database in 2021. Nucleic Acids Res 49: D412–D419

Nardini M (2003) CtBP/BARS: a dual-function protein involved in transcription co-repression and Golgi membrane fission. EMBO J 22: 3122–3130s

Necci M, Piovesan D, Clementel D, Dosztányi Z & Tosatto SCE (2021) MobiDB-lite 3.0: fast consensus annotation of intrinsic disorder flavors in proteins. Bioinformatics 36: 5533–5534

Nibu Y, Zhang H & Levine M (1998) Interaction of Short-Range Repressors with Drosophila CtBP in the Embryo. Science (80-) 280: 101–104

Nimura K, Ura K, Shiratori H, Ikawa M, Okabe M, Schwartz RJ & Kaneda Y (2009) A histone H3 lysine 36 trimethyltransferase links Nkx2-5 to Wolf–Hirschhorn syndrome. Nature 460: 287–291

Pfreundt U, James DP, Tweedie S, Wilson D, Teichmann SA & Adryan B (2010) FlyTF: improved annotation and enhanced functionality of the Drosophila transcription factor database. Nucleic Acids Res 38: D443–D447

Qi D, Bergman M, Aihara H, Nibu Y & Mannervik M (2008) Drosophila Ebi mediates Snail-dependent transcriptional repression through HDAC3-induced histone deacetylation. EMBO J 27: 898–909

Quinlan KGR, Verger A, Kwok A, Lee SHY, Perdomo J, Nardini M, Bolognesi M & Crossley M (2006) Role of the C-Terminal Binding Protein PXDLS Motif Binding Cleft in Protein Interactions and Transcriptional Repression. Mol Cell Biol 26: 8202–8213

Ramazzotti M, Monsellier E, Kamoun C, Degl’Innocenti D & Melki R (2012) Polyglutamine Repeats Are Associated to Specific Sequence Biases That Are Conserved among Eukaryotes. PLoS One 7: e30824

Ravarani CN, Erkina TY, De Baets G, Dudman DC, Erkine AM & Babu MM (2018) High-throughput discovery of functional disordered regions: investigation of transactivation domains. Mol Syst Biol 14

Reiter F, Wienerroither S & Stark A (2017) Combinatorial function of transcription factors and cofactors. Curr Opin Genet Dev 43: 73–81

Ryu JR & Arnosti DN (2003) Functional similarity of Knirps CtBP-dependent and CtBP-independent transcriptional repressor activities. Nucleic Acids Res 31: 4654–4662

Sabari BR, Dall’Agnese A, Boija A, Klein IA, Coffey EL, Shrinivas K, Abraham BJ, Hannett NM, Zamudio A V., Manteiga JC, et al (2018) Coactivator condensation at super-enhancers links phase separation and gene control. Science (80-) 361

Salichs E, Ledda A, Mularoni L, Albà MM & de la Luna S (2009) Genome-Wide Analysis of Histidine Repeats Reveals Their Role in the Localization of Human Proteins to the Nuclear Speckles Compartment. PLoS Genet 5: e1000397

Sanborn AL, Yeh BT, Feigerle JT, Hao C V., Townshend RJL, Aiden EL, Dror RO & Kornberg RD (2021) Simple biochemical features underlie transcriptional activation domain diversity and dynamic, fuzzy binding to mediator. Elife 10: 1–77

Schwanhäusser B, Busse D, Li N, Dittmar G, Schuchhardt J, Wolf J, Chen W & Selbach M (2011) Global quantification of mammalian gene expression control. Nature 473: 337–342

Shlyueva D, Stampfel G & Stark A (2014) Transcriptional enhancers: From properties to genome-wide predictions. Nat Rev Genet 15: 272–286

Smith ST & Jaynes JB (1996) A conserved region of engrailed, shared among all en-, gsc-, Nk1-, Nk2-and msh-class homeoproteins, mediates active transcriptional repression in vivo. Development 122: 3141–3150

Smyth GK (2004) Linear Models and Empirical Bayes Methods for Assessing Differential Expression in Microarray Experiments. Stat Appl Genet Mol Biol 3: 1–25

Soto LF, Li Z, Santoso CS, Berenson A, Ho I, Shen VX, Yuan S & Fuxman Bass JI (2022) Compendium of human transcription factor effector domains. Mol Cell 82: 514–526

Staller M V., Holehouse AS, Swain-Lenz D, Das RK, Pappu R V. & Cohen BA (2018) A High-Throughput Mutational Scan of an Intrinsically Disordered Acidic Transcriptional Activation Domain. Cell Syst 6: 444-455.e6

Staller M V., Ramirez E, Kotha SR, Holehouse AS, Pappu R V. & Cohen BA (2022) Directed mutational scanning reveals a balance between acidic and hydrophobic residues in strong human activation domains. Cell Syst 13: 334-345.e5

Stampfel G, Kazmar T, Frank O, Wienerroither S, Reiter F & Stark A (2015) Transcriptional regulators form diverse groups with context-dependent regulatory functions. Nature 528: 147–151

Stark A, Lin MF, Kheradpour P, Pedersen JS, Parts L, Carlson JW, Crosby MA, Rasmussen MD, Roy S, Deoras AN, et al (2007) Discovery of functional elements in 12 Drosophila genomes using evolutionary signatures. Nature 450: 219–232

Tanaka Y, Kawahashi K, Katagiri Z-I, Nakayama Y, Mahajan M & Kioussis D (2011) Dual Function of Histone H3 Lysine 36 Methyltransferase ASH1 in Regulation of Hox Gene Expression. PLoS One 6: e28171

Tapia-Ramírez J, Eggen BJL, Peral-Rubio MJ, Toledo-Aral JJ & Mandel G (1997) A single zinc finger motif in the silencing factor REST represses the neural-specific type II sodium channel promoter. Proc Natl Acad Sci 94: 1177–1182

Taus T, Köcher T, Pichler P, Paschke C, Schmidt A, Henrich C & Mechtler K (2011) Universal and Confident Phosphorylation Site Localization Using phosphoRS. J Proteome Res 10: 5354–5362

Tolkunova EN, Fujioka M, Kobayashi M, Deka D & Jaynes JB (1998) Two distinct types of repression domain in engrailed: one interacts with the groucho corepressor and is preferentially active on integrated target genes. Mol Cell Biol 18: 2804–14

Tycko J, DelRosso N, Hess GT, Aradhana, Banerjee A, Mukund A, Van M V., Ego BK, Yao D, Spees K, et al (2020) High-Throughput Discovery and Characterization of Human Transcriptional Effectors. Cell 183: 2020-2035.e16

Vaquerizas JM, Kummerfeld SK, Teichmann SA & Luscombe NM (2009) A census of human transcription factors: function, expression and evolution. Nat Rev Genet 10: 252–263

Waterhouse AM, Procter JB, Martin DMA, Clamp M & Barton GJ (2009) Jalview Version 2—a multiple sequence alignment editor and analysis workbench. Bioinformatics 25: 1189–1191

Wysocka J, Myers MP, Laherty CD, Eisenman RN & Herr W (2003) Human Sin3 deacetylase and trithorax-related Set1/Ash2 histone H3-K4 methyltransferase are tethered together selectively by the cell-proliferation factor HCF-1. Genes Dev 17: 896–911

Zabidi MA, Arnold CD, Schernhuber K, Pagani M, Rath M, Frank O & Stark A (2015) Enhancer-core-promoter specificity separates developmental and housekeeping gene regulation. Nature 518: 556–559

Zargar ZU & Tyagi S (2012) Role of Host Cell Factor-1 in cell cycle regulation. Transcription 3: 187–192

Zhang J-S, Moncrieffe MC, Kaczynski J, Ellenrieder V, Prendergast FG & Urrutia R (2001) A Conserved α-Helical Motif Mediates the Interaction of Sp1-Like Transcriptional Repressors with the Corepressor mSin3A. Mol Cell Biol 21: 5041–5049

## Preprint references

Emms DM, Kelly S. (2018) OrthoFinder: phylogenetic orthology inference for comparative genomics. bioRxiv 10.1101/267914 [PREPRINT]

